# Extracellular Vesicles Derived from Activated Dendritic Cells Loaded with Curcumin Promote Early Activation-associated Functional and Molecular Reprogramming of Primary CD8^+^ T Cells

**DOI:** 10.64898/2026.06.30.735629

**Authors:** Stefan Mihai Dragan, Laura Patras, Marta-Szilvia Meszaros, Ovidiu Ilie Pavel, Cristian V.A. Munteanu, Raluca Borlan, Monica Focsan, Alejandro Bernabeu Martinez, Ana Melero, Loredana Saveanu, Manuela Banciu, Alina Sesarman

## Abstract

Extracellular vesicles (EVs) derived from activated dendritic cells (DCs) are promising cell-free mediators capable of shaping CD8^+^ T-cell responses. However, their early molecular and functional effects on CD8^+^ T cells remain incompletely characterized, and whether engineering activated DC-derived EVs with immunomodulatory cargo can fine-tune these responses remains largely unexplored. Here, we investigated whether curcumin loading into EVs derived from CpG-activated and peptide-pulsed DC2.4 cells (EV-ACT) modulates early activation of primary CD8^+^ T cells. EVs were isolated by ultrafiltration coupled with size-exclusion chromatography (UF-SEC) and characterized physicochemically and molecularly. Exploratory proteomic profiling identified an activation-associated EV protein signature enriched in antigen-processing and immune-related pathways. Curcumin loading achieved an encapsulation efficiency of 16.4% while preserving EV properties, and spectral confocal fluorescence microscopy revealed heterogeneous fluorescence emission patterns consistent with distinct EV-associated curcumin microenvironments. Following rapid cellular association, EV-ACT promoted early CD8^+^ T-cell activation, inducing an effector-like phenotype characterized by increased CD69 expression, TNF-α and Granzyme B production, and reduced Bcl-2 levels without compromising cell viability. Unlike free curcumin, EV-mediated curcumin delivery selectively reinforced these immunostimulatory responses by significantly increasing CD69 expression and STAT3 phosphorylation, sustaining early activation-associated functional and molecular reprogramming of primary CD8^+^ T cells.

## Introduction

Extracellular vesicles (EVs) are lipid bilayer–enclosed nanoparticles released by most cell types that mediate intercellular communication through the transfer of bioactive cargo (El Andaloussi et al., 2013; Verma et al., 2015; Zhou et al., 2025). In cancer, EV-mediated communication within the tumor microenvironment (TME) plays a central role in regulating immune surveillance, immune evasion and therapeutic response (Buzas, 2023; Mahmoudi et al., 2023). Depending on their cellular origin, molecular cargo, and recipient-cell interactions, EVs can either suppress anti-tumor immunity (Huang et al., 2024; Patras et al., 2022) or promote anti-tumor immune responses by regulating immune-cell recruitment, antigen presentation and costimulatory signaling as well as by facilitating communication at the immune synapse (Céspedes et al., 2022; Kuang et al., 2025; Mittelbrunn et al., 2011; Ruiz-Navarro et al., 2024; Semionatto et al., 2020; Srivastava et al., 2021; Wu et al., 2020).

The immunological relevance of EVs was first demonstrated by Raposo and colleagues, who showed that B cell–derived EVs carry MHC class II–peptide complexes capable of modulating T-cell responses (Raposo et al., 1996). Since then, immune cell-derived EVs emerged as active regulators of adaptive immunity. Dendritic cell-derived EVs are particularly relevant because they retain key antigen-presenting features of their parental cells, including MHC class I and II molecules, costimulatory proteins, adhesion molecules (ICAMs) and tetraspanins, positioning them as promising cell-free mediators that can orchestrate antigen-specific immune responses through antigen dissemination, cross-dressing, immune-cell communication and amplification of CD8^+^ T-cell responses (Q. Liu et al., 2016; Moya-Guzmán et al., 2023; Pitt et al., 2016; W. Wang et al., 2025).

CD8^+^ T cells are central mediators of antitumor immunity through their direct capacity to recognize malignant cells and eliminate them via cytotoxic effector mechanisms, including pro-inflammatory cytokine secretion and granule-mediated killing (Koh et al., 2023; Yao et al., 2024). Previous studies have shown that dendritic cell-derived EVs carrying tumor-associated antigens can elicit antigen-specific T-cell responses, promote cytotoxic effector function and enhance anti-tumor immunity both *in vitro* and *in vivo* (Bu et al., 2015; Hay & Slansky, 2022; Veerman et al., 2023). More recently, engineering dendritic cell-derived EVs through modification of their surface molecules or molecular cargo has further enhanced cytotoxic T-cell responses and therapeutic efficacy (Bu et al., 2015; Chen et al., 2025; Fernández-Delgado et al., 2020; Zeng et al., 2026). However, these studies have largely evaluated downstream therapeutic endpoints, whereas the earliest molecular and functional events elicited in primary CD8 T cells following exposure to activated dendritic cell-derived EVs remain poorly understood. Early activation events are known to shape subsequent CD8^+^ T-cell differentiation and effector function (Obar & Lefrançois, 2010; Petkau et al., 2022). Emerging evidence also indicates that anti-tumor CD8^+^ T-cell responses are established through sequential activation programs, with early priming preceding acquisition of full effector function within the tumor (Prokhnevska et al., 2023). Therefore, characterizing these initial responses may guide the rational engineering of next-generation EV-based immunotherapies.

Identifying strategies capable of further potentiating these early activation events could improve the therapeutic efficacy of dendritic cell-derived EVs. Curcumin is a natural polyphenolic molecule extracted from the rhizome of *Curcuma longa*, with demonstrated pleiotropic, context- and dose-dependent biological effects including anti-inflammatory, anti-angiogenic and immunostimulatory properties (Ma et al., 2025; Zheng et al., 2025a). Curcumin has been reported to modulate CD8^+^ T-cell responses through mechanisms involving inflammatory signaling, cell survival and effector function. At low concentrations (10 µM), curcumin could prime immune responses by potentiating CD8^+^ T-cell activity through activation of NF-κB signaling and induction of anti-apoptotic programs (Guo et al., 2021; L. Liu et al., 2021a; Zheng et al., 2025a). However, the therapeutic potential of curcumin remains limited by poor aqueous solubility and low systemic bioavailability (Guzman-Villanueva et al., 2013). Beyond their physiological and pathological functions, EVs have attracted considerable interest as natural drug delivery systems owing to intrinsic biocompatibility, enhanced stability in biological fluids, capacity to bypass biological barriers, and ability to transport endogenous or exogenous cargo, offering several advantages over conventional synthetic nanocarriers (H. I. Kim et al., 2024). Loading curcumin into EVs or other nanocarrier systems has therefore emerged as a strategy to overcome these shortcomings (Aravind & Krishnan, 2016; Hasan et al., 2014; Sesarman et al., 2019).

Nevertheless, whether loading activated dendritic cell-derived EVs with an immunomodulatory cargo can further augment their intrinsic immunostimulatory properties and fine-tune the earliest molecular and functional responses of primary CD8^+^ T cells remains insufficiently explored. Therefore, in this proof-of-concept study, we comprehensively characterized EVs isolated from CpG-activated, TRP-2-pulsed murine DC2.4 cells for their physicochemical properties and molecular cargo, including their proteomic profiling, and we evaluated curcumin loading in EVs and uptake by primary CD8^+^ T cells. Finally, we investigated whether EV-associated curcumin potentiates the early activation-associated molecular and functional reprogramming of primary CD8^+^ T cells induced by activated dendritic cell-derived EVs.

## Materials and Methods

### 2.1 DC2.4 cell culture and primary CD8^+^ T-cell isolation

The murine dendritic cell line DC2.4 (Sigma-Aldrich, SCC142, St. Louis, MO, USA) was maintained in RPMI-1640 medium (BioWest, MS025D1008, Nuaillé, France) supplemented with 10% heat-inactivated fetal bovine serum (FBS; BioWest, S1600, Nuaillé, France), 100 IU/mL penicillin and 100 µg/mL streptomycin (Sigma-Aldrich, P4333, St. Louis, MO, USA), 4 mM L-glutamine (Lonza, BE17-605E, Verviers, Belgium), 10 mM sodium pyruvate (BioWest, L0642, Nuaillé, France), and 1x minimum essential medium (MEM) non-essential amino acids (NEAAs) without L-glutamine (BioWest, X0557, Nuaillé, France) (Lu et al., 2024).

Female C57BL/6 mice (6-8 weeks old) were obtained from the "Cantacuzino" National Medico-Military Institute for Research and Development (Bucharest, Romania) and maintained under standard housing conditions with *ad libitum* access to food and water. All animal procedures were approved by the Ethics Committee of Babes-Bolyai University (Reg. No. 15925/15.11.2022). Primary CD8^+^ T cells were isolated from pooled lymph nodes of 2–3 female C57BL/6 mice (8–10 weeks old) using the MojoSort™ Mouse CD8 T Cell Isolation Kit (BioLegend, 480035, San Diego, CA, USA). Briefly, lymph nodes were mechanically dissociated, filtered through a 40 μm cell strainer into sterile PBS (BioWest, L0616, Nuaillé, France) containing 0.2 mM EDTA and 1% bovine serum albumin (BSA). After washing, cells were resuspended in PBS containing 1% BSA and subjected to magnetic negative selection using a cocktail of biotinylated antibodies against non-CD3^+^CD8^+^ immune cell populations according to the manufacturer’s instructions. Cell viability was assessed by Trypan Blue exclusion and CD8^+^ T-cell purity was measured by flow cytometry. Purified CD8^+^ T cells were cultured in complete RPMI-1640 medium supplemented with 10% heat-inactivated FBS, 100 IU/mL penicillin, 100 μg/mL streptomycin, 4 mM L-glutamine, 10 mM sodium pyruvate, 1× MEM NEAAs, and 5 ng/mL recombinant murine Interleukin-2 (BioLegend, 575404, San Diego, CA, USA). Purified CD8^+^ T cells were allowed to recover for 3 h under standard culture conditions (37°C, 5% CO_2_) before subsequent experiments (Gao et al., 2022).

### 2.2 Generation and isolation of dendritic cell-derived EVs

DC2.4 murine dendritic cells were seeded in 10-cm tissue culture dishes in complete RPMI-1640 medium and cultured for 24 h. Cells were monitored using an inverted microscope (Axio Vert 1, Carl Zeiss, Oberkochen, Germany). At approximately 60% confluence, cells were divided into experimental groups. For activation, the culture medium was replaced with complete RPMI-1640 supplemented with 0.5 µM CpG (InvivoGen, TLRL-1826-5, Toulouse, France), an established *in vitro* dose for robust dendritic cell activation (Warren et al., 2000; Yazdani et al., 2021), together with 0.5 µM TRP-2 (180-188) peptide (AnaSpec, AS-61058, Fremont, CA, USA), to promote antigen loading and presentation (H.-M. Liu et al., 2003). Tyrosinase-related protein 2 (TRP-2) peptide was selected as the model tumor-associated antigen because vaccination with this peptide has demonstrated anti-melanoma activity in preclinical studies. Moreover, this epitope is presented by both murine MHC-I H-2Kb and human HLA-A2 molecules, supporting its translational relevance. CpG oligodeoxynucleotides (CpG-ODNs) were used to create a sufficient “danger signal” for DCs to mature and acquire a pro-inflammatory Th1 phenotype (Jordan et al., 2012; Mansour et al., 2007; Z. Xu et al., 2013). Control cells were maintained under identical culture conditions in complete RPMI-1640 without CpG or TRP-2 peptide.

Following 16 h of incubation, cells were detached using 0.5 mM EDTA in PBS, counted and reseeded at 4x10^6^ cells per 20-cm tissue culture dish in complete RPMI-1640 supplemented with 2% exosome-depleted FBS (Gibco, A2720801, Waltham, MA, USA). After 48 h, conditioned media from activated and non-activated cultures was collected for EV isolation. EVs isolated from activated and non-activated dendritic cells are hereafter referred to as EV-ACT and EV-NA, respectively.

EVs were isolated using ultrafiltration coupled with size-exclusion chromatography (UF-SEC) as previously described (Patras et al., 2022), to ensure reliable and efficient enrichment of small-sized EVs for further functional studies. Briefly, conditioned media were sequentially centrifuged at 500×g for 10 min and 3000×g for 20 min to remove cells and cellular debris, followed by filtration through a 0.22 μm membrane. Samples were concentrated to 1 mL by ultrafiltration using Vivaspin® 20 centrifugal units (100 kDa MWCO) (Sartorius, VS2041, Göttingen, Germany), before separation at 4°C on a Sephacryl S-200 HR column (Sigma, SLBW0078, St. Louis, MO, USA) equilibrated with sterile PBS and connected to an automated fraction collector connected to a temporizer and a peristaltic pump (Jung et al., 2021). Fractions corresponding to the previously established EV-containing peak at 280 nm (**Supplementary Figure S1**) were pooled and concentrated to a final volume of 500 μL. Protein concentration of EV-ACT and EV-NA preparations was determined using the Pierce™ BCA Protein Assay Kit (Thermo Scientific, 23225, Waltham, MA, USA) according to the manufacturer’s instructions.

### 2.3 Curcumin loading into EVs

To load curcumin into activated dendritic cell-derived EVs (EV-CURC), EV-ACT preparations were incubated with curcumin dissolved in 100% DMSO at a ratio of 1 mg curcumin per mg EV protein for 2 h at 37°C under constant agitation (400 rpm). Following incubation, preparations were centrifuged at 1000*xg* for 5 min to remove aggregates. Free curcumin was subsequently removed at room temperature by size-exclusion chromatography (SEC) using a Sephadex G-25 column (Sigma-Aldrich, S-5772, St. Louis, MO, USA) equilibrated with PBS as the mobile phase (Ó’Fágáin et al., 2017). EV-CURC-containing fractions were pooled, concentrated by ultrafiltration to a final volume of 250 µL. Encapsulation efficiency of curcumin was determined fluorometrically. Briefly, 5 µL of EV-CURC were diluted 30-fold in absolute ethanol to disrupt EV membranes and extract EV-associated curcumin. Following centrifugation (1000x*g*, 5 min), fluorescence of the supernatant was measured using a FLUOstar Omega microplate reader (BMG Labtech, Ortenberg, Germany) (Ex 485 nm/Em 520 nm) against a standard curve prepared from curcumin serial dilutions in absolute ethanol ranging from 0.0875 to 3.5 µg/mL. Encapsulation efficiency (%) was calculated as the amount of EV-associated curcumin relative to the total amount of curcumin initially added × 100.

### 2.4 Physicochemical characterization and stability analysis of EVs

Hydrodynamic diameter, polydispersity index (PDI), and zeta potential of EV-ACT and EV-CURC were determined by dynamic light scattering (DLS) using a Zetasizer Nano ZS90 (Malvern Panalytical Ltd., UK) as previously described (Holca et al., 2025). Measurements were performed in disposable square polystyrene cuvettes (DTS0012). The physicochemical stability of EV-CURC was evaluated under biologically relevant conditions. A pH-dependent stability study was performed at pH 7.4, 6.2, and 5.0, corresponding to physiological, mildly acidic, and endosomal environments, respectively, at 25°C and 37°C. In parallel, a time-dependent stability study was conducted at the same pH values immediately after preparation (day 0) and following 15 and 30 days of storage at 25°C. Between measurements, suspensions were stored at 4°C in the dark to maintain vesicle integrity. All measurements were performed in triplicate.

Particle size distribution and concentration of representative EV-ACT preparations were further characterized by nanoparticle tracking analysis (NTA) using a NanoSight LM10 instrument (Malvern Instruments Ltd., Malvern, UK) equipped with a 405 nm laser and an sCMOS camera. Data were acquired and analyzed using NTA software version 3.3 (Dev Build 3.3.104). Detection threshold (5), camera gain (15), and blur settings were automatically adjusted by the software. For each sample, three 30-second videos were acquired at a frame rate of 30 fps while maintaining a constant sample temperature throughout acquisition. Prior to analysis, samples were diluted in 0.22 μm-filtered PBS to obtain particle concentrations within the manufacturer’s recommended range (20–120 particles per frame) (Filipe et al., 2010).

### 2.5 Flow cytometric characterization of dendritic cells and CD8^+^ T cells

Flow cytometry was used to characterize dendritic cell activation, determine CD8^+^ T-cell purity following isolation, and quantify early T-cell activation. To confirm dendritic cell activation, the expression of CD86, MHC class II, TRP-2 and MHC class I was evaluated in non-activated (NA DC2.4) and CpG/TRP-2-treated (ACT DC2.4) cells by flow cytometry. Cells were detached using 0.5 mM EDTA in PBS, and 2x10^5^ cells were washed by centrifugation at 350x*g* for 5 min with staining buffer (PBS containing 1% BSA), and incubated with the Fc receptor blocking reagent (TruStain FcX™, 101320, San Diego, CA, USA) according to the manufacturer’s instructions. Cells were stained with PE-conjugated anti-CD86 and PE-Cy5-conjugated anti-MHC class II antibodies for 20 min, or with primary antibodies against TRP-2 and MHC class I for 1 h followed by the corresponding Alexa Fluor 532- and PE-Cy5-conjugated secondary antibodies for 20 min (**Supplementary Table S1**). All incubations were performed at room temperature, in the dark. Following staining, cells were washed twice with staining buffer, incubated with DRAQ7 viability dye (Invitrogen, D15105, Waltham, MA, USA) according to the manufacturer’s instructions before acquisition. The corresponding gating strategies are presented in **Supplementary Figures S2-S6**.

Primary CD8^+^ T-cell purity following magnetic isolation was assessed by flow cytometry using a PE-conjugated anti-mouse CD8 antibody (**Supplementary Table S1**). To assess early CD8^+^ T-cell activation, 2x10^5^ cells were washed twice with staining buffer by centrifugation at 500x*g* for 5 min, incubated with Fc receptor blocking reagent and stained with PE-conjugated anti-mouse CD69 antibody (**Supplementary Table S1**) for 20 min at room temperature, in the dark. Following two washes with staining buffer, DRAQ7 viability dye was added according to the manufacturer’s instructions before flow cytometric acquisition. The corresponding gating strategies are presented in **Supplementary Figures S7 and S8**.

Flow cytometry was performed using a Guava® Muse™ Cell Analyzer (Luminex, Austin, TX, USA) and data were analyzed using the integrated Guava® Muse® software (Sheen et al., 2017). Antibodies used are listed in **Supplementary Table S1**.

### 2.6 Western blot analyses

Western blot analysis was performed to characterize EV preparations and their parental DC2.4 cells, validate selected proteomic findings, and investigate early molecular changes induced in primary CD8^+^ T cells. EVs, DC2.4 cells and CD8^+^ T cells were lysed in RIPA buffer (Pierce, Thermo Fisher Scientific, XK356640, Waltham, MA, USA) supplemented with phosphatase (Roche, 04906845001, Basel, Switzerland) and protease inhibitors (Roche, 11697498001, Basel, Switzerland). Following centrifugation at 15,000×g, protein lysates were collected and protein concentration was determined using the Pierce™ BCA Protein Assay Kit (Thermo Scientific, 23225, Waltham, MA, USA).

For EVs and parental cell characterization, 30 µg of total protein were resolved by 10% SDS-PAGE, transferred to PVDF membranes and probed with antibodies against Tumor Susceptibility Gene 101 (TSG101), Stomatin (STOM), Galectin-3-binding protein (LGALS3BP) and β-actin (ACTB) as a reference protein, while Golgi matrix protein (GM130) was included as a negative EV marker to assess cellular contamination of EV preparations, in accordance with the MISEV recommendations (Théry et al., 2018; Welsh et al., 2024). (**Supplementary Table S1**). Heat shock protein 70 (HSP70) and fatty acid synthase (FASN) were analyzed to validate selected proteomic findings. For analysis of primary CD8^+^ T-cell lysates, 2.5 µg of total protein were loaded per lane onto 4–20% Mini-PROTEAN® TGX™ Precast Protein Gels (Bio-Rad, 4561094, Hercules, CA, USA) to evaluate the expression levels of phosphorylated Signal Transducer and Activator of Transcription 3 (p-STAT3), phosphorylated Nuclear Factor kappa-light-chain-enhancer of activated B cells (p-NF-κB), Poly(ADP-ribose) polymerase family member 14 (PARP14) and B-cell lymphoma 2 (Bcl-2).

Primary and HRP-conjugated secondary antibodies were diluted in 5% BSA prepared in Tris-buffered saline containing 0.1% Tween-20, according to **Supplementary Table S1**. Membranes were incubated with primary antibodies overnight at 4°C and with secondary antibodies for 1 h at room temperature. Immunoreactive bands were detected using ECL Clarity (Bio-Rad, 170-50611, Hercules, CA, USA) for EV and parental cell characterization, whereas Clarity Max™ Western ECL Substrate (Bio-Rad, 1705062, Hercules, CA, USA) was used for the analysis of CD8^+^ T cells. Images were acquired with a ChemiDoc™ Touch Imaging System (Bio-Rad, Hercules, CA, USA) and analyzed using Image Lab software version 6.1 (Bio-Rad, Hercules, CA, USA).

### 2.7 LC–MS/MS proteomic profiling of EVs

#### SDS-PAGE preparation and in-gel digestion

EV samples (EV-NA and EV-ACT) obtained from three independent biological preparations were incubated with SDS-containing Laemmli buffer in the presence of 8% β-mercaptoethanol for 5 minutes at 95°C. Samples (80 µg/lane) were resolved on 10% SDS-polyacrylamide gels and stained with Coomassie Brilliant Blue (Merck, 115444, Darmstadt, Germany). Each lane was subsequently excised into seven gel slices, and corresponding gel slices from each experimental condition were pooled in acetonitrile (99.5% ACS grade) in 1.5 mL tubes.

EV samples were subjected to in-gel digestion with trypsin according to a previously published protocol (Petrareanu et al., 2014; Popa et al., 2016). Briefly, following destaining with 50% acetonitrile/25 mM ammonium bicarbonate, each gel slice underwent disulfide bond reduction with DTT, alkylation with 55 mM Iodocetamide, and overnight trypsin digestion at 37°C. The resulting peptides were extracted twice using 50% acetonitrile containing 5% formic acid and once using 95% acetonitrile, concentrated to dryness in a SpeedVac and stored at -20°C until LC–MS/MS analysis.

#### LC-MS/MS analysis

Peptides derived from EV-NA and EV-ACT samples were reconstituted in solvent A (0.06% formic acid (FA) and 2% acetonitrile (ACN)) and analyzed by LC-MS/MS using an Easy-nanoLC 1200 system connected online to an LTQ-Orbitrap Velos Pro mass spectrometer (Thermo Fisher Scientific, Bremen, Germany), using acquisition parameters similar to those previously described (Munteanu et al., 2021). Peptides were first online desalted using a trap column connected directly to the analytical column used for separation and to the waste side through a liquid junction cross. Peptides were separated on a reversed-phase C18 Acclaim PepMap 100 column using a 2-30% solvent B (0.06% FA and 80% ACN) gradient for 90 minutes, followed by a rapid increase to 90% solvent B for 5 minutes. The eluted peptides were detected using a data dependent acquisition strategy under which the top five most abundant peptide ions from the MS1 scan acquired using the Orbitrap mass analyzer were fragmented either in the linear ion trap to generate collision-induced dissociation (CID) scans or in the higher-energy collisional dissociation (HCD) cell with Orbitrap detection. Multiple injections were performed from the same vial and HCD and CID datasets were obtained in separate analytical runs. The lock mass was enabled for the MS1 acquisition of scans using the m/z value of 445.120025. Dynamic exclusion was activated with an exclusion duration of 30 s and an exclusion list size of 500, using an exclusion mass width relative low and high limits to 10 ppm.

#### Raw data analysis

The acquired raw data were searched against the fasta sequences of the *Mus musculus* UniProtKB reference proteome (UniProtKB 17,213 sequences as of 02.2025) using the SequestHT algorithm integrated into Proteome Dicoverer v2.5, selecting trypsin as the digestion enzyme and a maximum of two missed cleavages. Carbamidomethylation of cysteine residues was specified as a static modification and methionine oxidation was considered a dynamic modification. For precursor ions the maximum allowed mass deviation was 10 ppm, while for the fragment ions 0.02 Da for scans acquired in the Orbitrap analyzer and 0.5 Da for the scans acquired in the ion trap. False Discovery Rate (FDR) estimation of the same data was searched against a decoy database, and the resulting peptide-spectrum matches (PSM) were further filtered to 1% FDR using the default values in the consensus step for peptide and protein validation (Elias & Gygi, 2007). Raw LC–MS/MS files and processed proteomic datasets were deposited in the MassIVE repository under accession **MSV000101874**.

#### Bioinformatic analysis and data visualization

Bioinformatic analyses were performed in R version 4.4.2 (R Foundation for Statistical Computing, Vienna, Austria) using the tidyverse package (Wickham et al., 2019). Protein abundances were log_2_-transformed, and log_2_ fold changes (log_2_FC) between EV-NA and EV-ACT were calculated. Proteins exhibiting an absolute |log_2_FC| > 1 were selected for exploratory functional enrichment analyses using clusterProfiler and gprofiler2 against Gene Ontology (GO) and KEGG databases with org.Mm.eg.db for mouse gene annotation. Data visualization was performed using ggplot2, enrichplot, and pathview. Differentially abundant proteins were compared with the top 100 EV proteins listed in the ExoCarta database (Keerthikumar et al., 2016). Overlapping proteins were visualized as log_10_-transformed heatmaps using ComplexHeatmap (Gu et al., 2016), and GO molecular function and biological process enrichment analyses were performed. For a manually curated panel of proteins involved in MHC-I antigen presentation, proteasome/immunoproteasome function, ER protein folding, antigen processing, EV biogenesis and trafficking, and immune adhesion, protein abundances were transformed using log_2_(abundance+1), and heatmaps were generated with pheatmap without hierarchical clustering to preserve predefined biological functional groupings. All analyses were performed independently using both HCD- and CID-derived protein abundance datasets, as both yielded comparable biological trends. Only HCD-based analyses were presented throughout the manuscript for clarity and because this this fragmentation method was reported to provide higher peptide identification rates and more confident peptide-spectrum matches than conventional CID for tryptic proteomic samples (Frese et al., 2011). Illustrations and experimental workflow figures were created using BioRender (BioRender.com).

### 2.8 Spectrofluorometric characterization of EV-associated curcumin

Fluorescence emission were recorded using a Jasco FP-6500 spectrofluorometer (Jasco International Co., Ltd., Tokyo, Japan). Measurements were performed in 5 × 5 mm quartz cuvettes (Hellma, Germany) with excitation wavelengths set to 430, 450, and 470 nm, as described in the literature (Radha et al., 2024). Emission spectra were acquired in the 480–800 nm range, using identical slit widths and integration times for all samples. Measurements of spectral intensity were carried out on free curcumin (free CURC) in PBS, EV-ACT, and EV-CURC under identical experimental conditions to enable direct comparison. Because free CURC does not emit fluorescence in aqueous solution, an equivalent concentration of curcumin dissolved in absolute ethanol was analyzed under identical acquisition settings as a reference, to evaluate fluorescence stability and spectral behavior of the compound.

### 2.9 Spectral confocal fluorescence microscopy of EV-associated curcumin

Confocal spectral fluorescence images of EV-CURC deposited directly onto glass microscopy slides were acquired using a ZEISS LSM 900 confocal microscope (Zeiss, Oberkochen, Germany) mounted on an Axio Observer Z1/7 inverted stand (Zeiss, Oberkochen, Germany) and equipped with a Plan-Apochromat 63×/1.40 Oil DIC M27 objective. Samples were excited using a 405 nm diode laser operated at 0.1% power. Fluorescence emission spectra were collected over the 450–644 nm spectral range in 13 sequential detection windows with a channel width of 15 nm. Representative images were displayed for λ_em = 457, 472, 502, 517, and 532 nm. Images were acquired in frame-scan mode with a scan area of 2.0, a pixel size of 0.059 μm, and an image resolution of 287 × 287 pixels. Image acquisition and spectral separation were performed using the ZEISS ZEN software (Storti et al., 2023).

### 2.10 Cellular uptake of EV-CURC by primary CD8^+^ T cells

To evaluate the cellular uptake of EV-CURC, 1x10^5^ purified primary CD8^+^ T cells were seeded onto Ibidi glass-bottom µ-Dish 35 mm (ibidi GmbH, 81156, Gräfelfing, Germany) and allowed to rest for 3 h in complete RPMI-1640 medium under standard culture conditions, as described above. Cells were subsequently incubated with 5 µM EV-CURC or free CURC for 30 min, while an equivalent volume of PBS corresponding to the EV suspension volume was used as negative control. After incubation, cells were washed twice with sterile PBS by centrifugation (500 × g) to remove unbound EV-CURC or free CURC, fixed in 4% paraformaldehyde (PFA) for 10 minutes at 4°C, washed twice with ice-cold PBS, and stained with 1 µg/mL DAPI (4′,6-diamidino-2-phenylindole, Sigma-Aldrich, D9542, St. Louis, MO, USA), for 10 minutes at room temperature protected from light. Fluorescence images were acquired using a Nikon Eclipse Ti2-E inverted fluorescence microscope (Nikon Instruments Inc., Tokyo, Japan), operated with NIS-Elements software (version 5.12) for microscope control, image acquisition and processing (Holca et al., 2025).

### 2.11 Primary CD8^+^ T cell-stimulation and treatment

To evaluate the effects of EV-delivered curcumin on primary CD8**^+^**T-cell activation and early functional responses, purified CD8**^+^** T cells were treated with EV-CURC for 12 or 24 h at concentrations equivalent to 1 µM and 5 µM free curcumin (CURC). Control EV preparations (EV-ACT) were added at protein concentrations equivalent to those present in the corresponding EV-CURC formulations. Untreated cells receiving an equivalent volume of PBS as that introduced with the EV-based treatments were considered as controls (untreated cells). CD8**^+^** T cells chemically stimulated with 81 nM phorbol 12-myristate 13-acetate (PMA) and 1.34 µM ionomycin (eBioscience™, 00-4970-93, Waltham, MA, USA) known to induce a robust antigen-independent pharmacological activation of T cells (Abdulhaqov et al., 2026), was considered as positive control throughout the manuscript and was used to validate subsequent measurements.

### 2.12 Enzyme-Linked Immunosorbent Assay (ELISA)

To concentration of murine Granzyme B released by primary CD8^+^ T cells into the culture was quantified using a commercial sandwich ELISA kit (Assay Genie, MOFI00260, Dublin, Ireland) according to the manufacturer’s instructions. Briefly, cell culture supernatant and recombinant murine Granzyme B standards (1.563-100 pg/mL) were added to antibody-precoated 96-well plates for 90 min at 37°C, followed by sequential incubation with biotinylated detection anti-mouse Granzyme B antibodies (60 min at 37°C), streptavidin-biotin complex (SABC) solution (30 min at 37°C) and tetramethylbenzidine (TMB) substrate at 37°C until a specific blue color appeared on the most concentrated standard. The reaction was stopped using the supplied stop solution and absorbance was measured at 450 nm using a FLUOstar Omega microplate reader (BMG Labtech, Ortenberg, Germany) (Lin et al., 2014).

The concentration of murine TNF-α released by primary CD8^+^ T cells into the culture was quantified using a commercial sandwich ELISA Development Kit (PeproTech, 315-01A-54, Cranbury, NJ, USA) according to the manufacturer’s instructions. Briefly, high-binding 96-well plates were coated overnight at room temperature with capture antibody (0.5 µg/mL) diluted in PBS. Plates were washed and blocked with 1% BSA for 1 h at room temperature. Cell culture supernatant and equal volumes of recombinant murine TNF-α standards (8–2000 pg/mL) were added in duplicate and incubated for 2 h at room temperature. After washing, biotinylated detection antibody (0.125 µg/mL) was added and incubated for 2 h, followed by incubation with streptavidin-HRP conjugate (0.025 µg/mL) for 30 min. Color development was achieved by incubating with TMB substrate for 20 min and the reaction was stopped with 1M HCl solution. Absorbance was measured at 450 nm with wavelength correction at 620 nm using a FLUOstar Omega microplate reader (BMG Labtech, Ortenberg, Germany) (Khakhum et al., 2021).

### 2.13 Statistical analysis

Statistical analysis was performed using GraphPad Prism version 11.0.0 (GraphPad Software, Boston, MA, USA). Differences between two groups were assessed using an unpaired Student’s *t* test, whereas comparisons among multiple groups were performed using one-way analysis of variance (ANOVA) followed by Dunnett’s multiple comparisons test for comparisons with the control group or Sidak’s multiple comparisons test for selected pairwise comparisons. Results are presented as mean ± standard deviation (S.D.). Statistical significance was defined as *p* < 0.05.

## 3. Results

### 3.1 Activated dendritic cells release extracellular vesicles with characteristic physicochemical and molecular features

Activation of DC2.4 cells was confirmed by the increased expression of CD86, CD80, MHC-I and TRP-2, whereas MHC class II expression remained unchanged (**Figure 1A**). Following activation, EVs were isolated from CpG-activated and TRP-2 peptide-pulsed DC2.4 cells according to the workflow shown in **Figure 1B**. DLS analysis revealed two particle populations, with a predominant population (83%) displaying a hydrodynamic diameter of approximately 160 nm, consistent with the expected size range of small EVs (Welsh et al., 2024), and a minor population (16%) with a diameter of approximately 11 nm (**Figure 1C**). EV preparations exhibited a PDI of 0.7 and a slightly negative zeta potential of -7.8 ± 0.2 mV (**Figure 1C**). NTA further confirmed the presence of a typical EV population, with a median particle diameter of 192 nm and a particle concentration of 1.66 x 10^12^ particles/mL (**Figure 1D**). Western blot analysis confirmed the presence of EV-associated protein markers, including TSG101, Stomatin, and LGALS3BP, whereas the Golgi marker GM130 was not detected in EV preparations (**Figure 1E**) (Matamoros Angles et al., 2024). Collectively, these findings demonstrate that the isolated EV preparations fulfilled the physicochemical and protein marker characterization criteria recommended by the MISEV2023 guidelines (Welsh et al., 2024).

**Figure 1.**
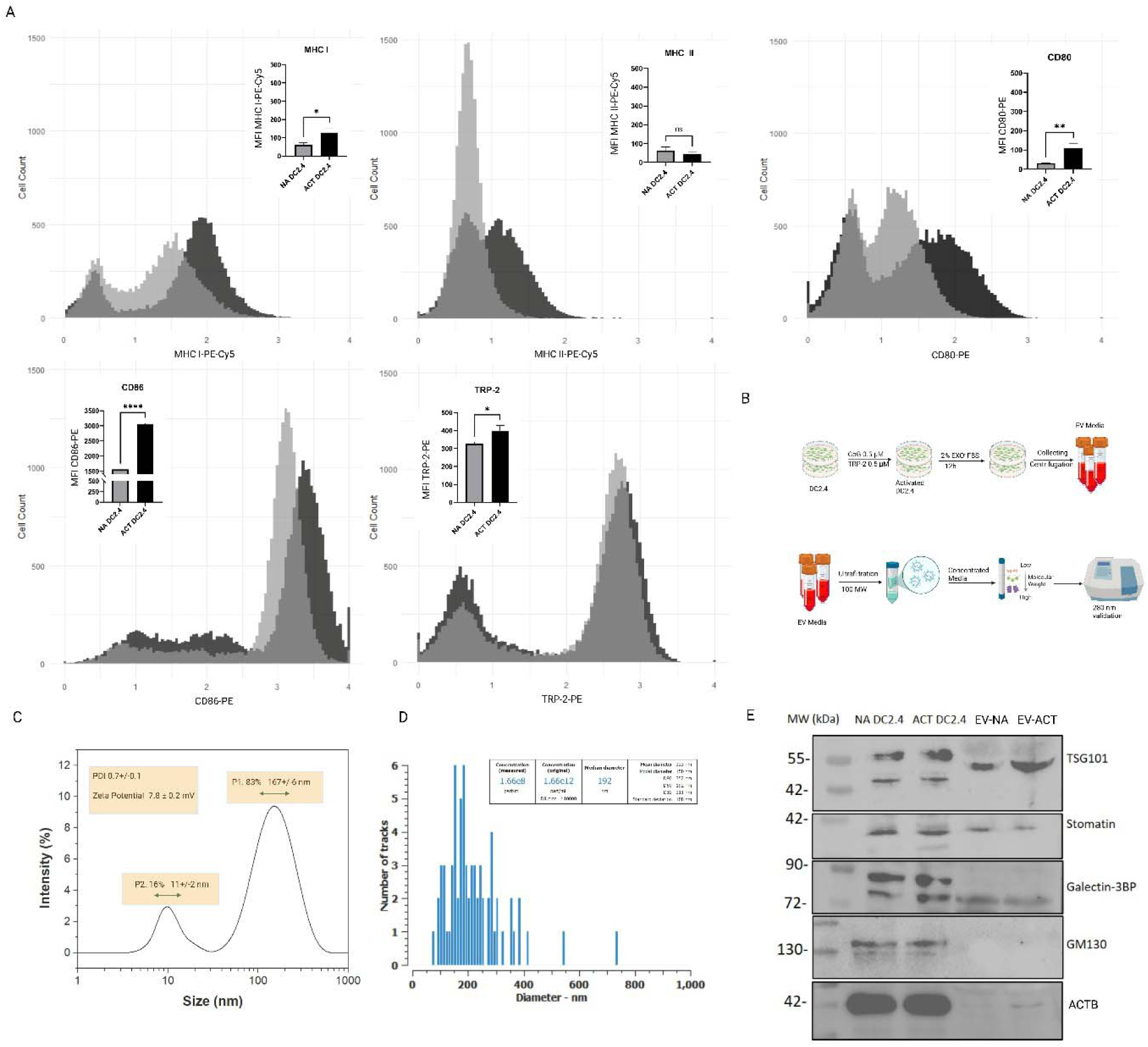
Isolation and characterization of extracellular vesicles derived from CpG-activated and TRP-2 peptide-pulsed dendritic cells (DC2.4). **A.** Flow cytometric analysis of dendritic cell activation following CpG stimulation and TRP-2 peptide-pulsing (mean ± S.D., n = 3 technical replicates). Unpaired Student’s t test was used to measure statistical significance; ns – not significant; P > .05; *, P < .05; **, P< .01; ***, P < .001. **B.** Schematic overview of the EV isolation workflow. **C.** Dynamic light scattering (DLS) analysis of EVs isolated from activated dendritic cells showing particle size distribution, polydispersity index (PDI), and zeta potential (n = 3 technical replicates). **D.** Representative nanoparticle tracking analysis (NTA) of EV-ACT. **E.** Western blot characterization of EV-associated proteins and the negative EV marker GM130. NA DC2.4 = non-activated murine dendritic cells; ACT DC2.4 = CpG-activated and TRP-2 peptide-pulsed murine dendritic cells; EV-NA = extracellular vesicles isolated from NA DC2.4 cells; EV-ACT = extracellular vesicles isolated from ACT DC2.4 cells.

### 3.2 Activated dendritic cell-derived EVs display an immunologically relevant proteomic cargo

LC–MS/MS proteomic profiling identified 1005 proteins in EV-NA and 1075 proteins in EV-ACT, demonstrating extensive proteomic coverage of both EV preparations. Comparison with EV-dedicated databases such as ExoCarta and EVpedia (D. Kim et al., 2013) revealed that 57% of proteins identified in EV-NA and 53% of proteins identified in EV-ACT overlapped with previously reported murine EV protein signatures from the queried databases, as shown in the Venn diagram (**Figure 2A**). Approximately 10% of the proteins identified in EV-NA and 11% in EV-ACT were not represented in either of the EV-dedicated databases (**Supplementary Table S2**), thereby suggesting that these proteins may represent previously unreported EV-associated proteins or context-dependent cargo related to the activation or culture (i.e., metabolic stress) conditions used in this study (Abramowicz et al., 2019).

**Figure 2.**
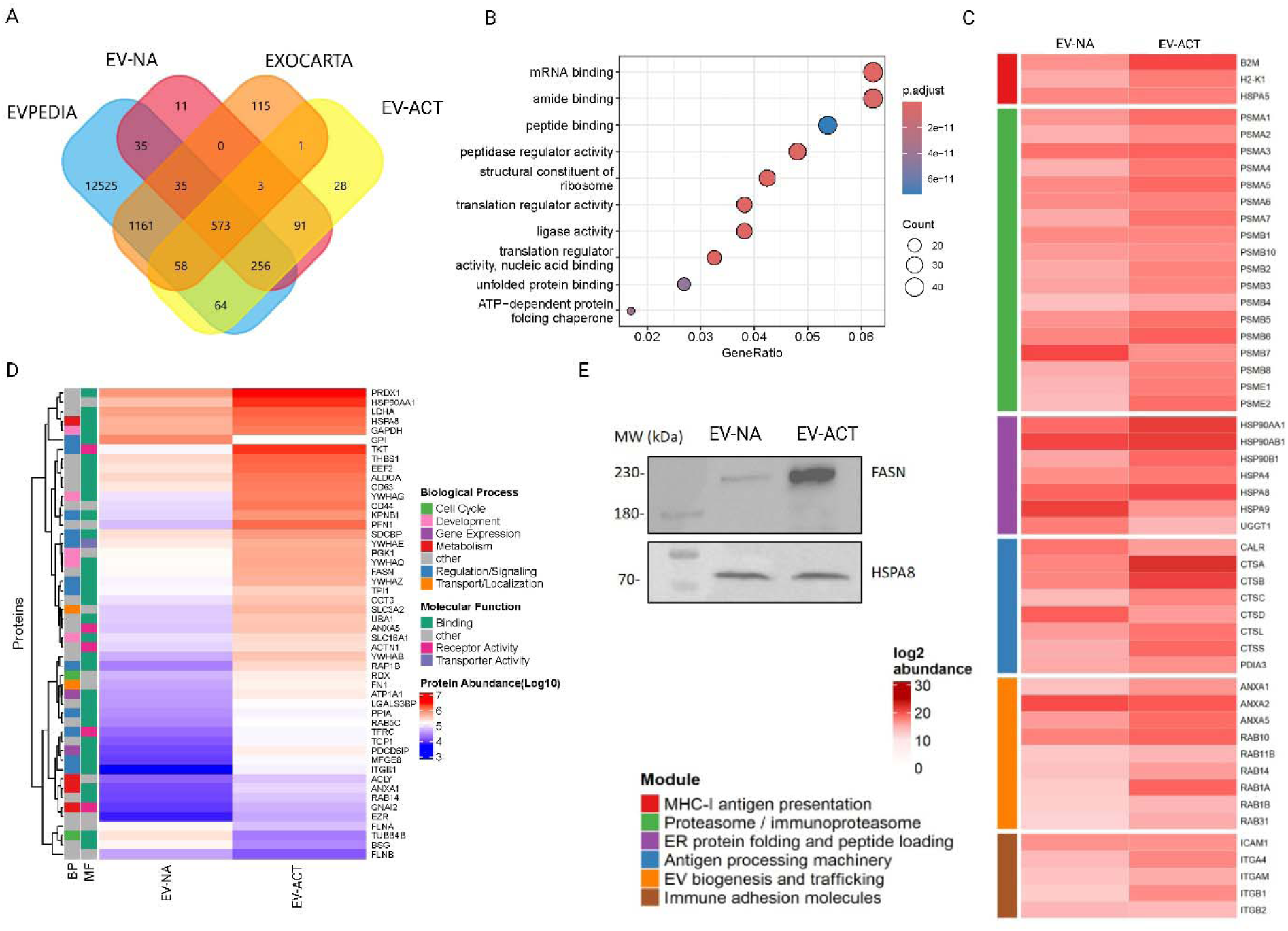
Proteomic characterization of EV-associated proteins derived from non-activated and activated dendritic cells, identified by mass spectrometry. **A.** Venn diagram showing the overlap between proteins identified in EV-NA and EV-ACT preparations and previously reported murine EV proteins curated in the ExoCarta and EVpedia databases. EV-NA = EVs isolated from non-activated DC2.4 cells; EV-ACT = EVs isolated from CpG-activated and TRP-2 peptide-pulsed DC2.4 cells. **B.** Gene Ontology enrichment analysis of differentially abundant proteins (DAPs) in EV-ACT compared to EV-NA; analysis was performed with R Studio and shows enrichment molecular function. **C**. Heatmap for differentially abundant proteins (relative abundance, log_2_ scale) involved in MHC-I antigen presentation, Proteasome and Immunoproteasome, ER protein folding, Antigen processing, EV biogenesis and trafficking, and Immune adhesion molecules. **D.** Heatmap of the 50 differentially abundant proteins (50) overlapping with the Top 100 most frequently reported EV proteins from ExoCarta, displayed as log_10_ protein abundance, together with the corresponding Gene Ontology enrichment analysis for these proteins: BP =biological process; MF = molecular function. **E.** Validation of selected proteins FASN and HSPA8 expression levels in EV-ACT versus EV-NA by western blot analysis.

To identify activation-associated EV proteins and to evaluate the enrichment of immune-related biological pathways, comparative proteomic analyses identified 655 proteins with differential abundance in EV-ACT (|log_2_FC| > 1, EV-ACT/EV-NA), demonstrating extensive remodeling of the EV proteome following dendritic cell activation. Exploratory Gene Ontology enrichment analysis of the EV-ACT-enriched proteins compared to EV-NA demonstrated an overrepresentation of proteins involved in mRNA binding and amide-binding, suggesting that EV-ACT may carry proteins associated with post-transcriptional regulation (Mateescu et al., 2017) and biosynthetic pathways relevant to immune cell activation (Buck et al., 2014) (**Figure 2B**). Among the enriched proteins, several were reported to actively participate in amino acid metabolism (NARS1, VARS1, RPLP2), as well as immune modulation through molecular chaperone activity (HSP90AA1, HSP70 and HSP90B1), cytoskeletal organization (ACTN1, AP2N1, and RAB1A) and antigen processing and presentation pathways (HSP70, HSP90A1 and MHC-I) **(Supplementary Figure S9)**. Notably, the proteins involved in antigen processing and presentation processes were preferentially enriched in EV-ACT, suggesting a potential role for these vesicles in antigen cross-presentation (**Figure 2C**) (Buzas, 2023) and subsequent activation of cytotoxic T cells (Curtsinger et al., 1999). Similarly, proteins associated with proteasome and immunoproteasome complexes (PSMA 1,3,5,7 and PSMB 5,6), as well as with endoplasmic reticulum (ER) processing pathways (HSP90AA1, HSP90AB1 and HSPA8) were also enriched in EV-ACT (**Figure 2C**, **Supplementary Figures S10 and S11)**, further supporting an activated antigen-processing phenotype.

To evaluate the EV identity of the activation-associated proteomic signature, differentially abundant proteins (DAPs) in EV-ACT compared to EV-NA were intersected with the top 100 EV proteins reported in the ExoCarta (**Figure 2D**). The overlapping proteins (approximately 50%) were further annotated with Gene Ontology enrichment analysis to show enriched molecular function and biological processes. Results showed that 90% of the proteins have higher abundance in EV-ACT, and are either canonical EV proteins such as CD63, LGALS3BP and HSPA8, or are involved in regulation/signaling (SDCBP, YWHAE, YWHAZ, TPI1), binding and receptor activity (TKT, ANXA5, ACTN1). Among these enriched proteins, FASN and HSPA8 were selected for validation by western blot (**Figure 2E**), due to their known association with EV biology and their roles in lipid metabolism (FASN) (Qian et al., 2018) and protein homeostasis, protein folding and EV cargo sorting (HSPA8) (Théry et al., 1999).

### 3.3 Curcumin loading preserves EV stability while generating distinct EV-associated physicochemical microenvironments

To exploit the intrinsic cargo delivery capacity of activated dendritic cell-derived EVs, curcumin (CURC), a pleiotropic immunomodulatory compound, was incorporated into EV-ACT by passive loading (Y. Wang et al., 2020) (**Figure 3A**). An encapsulation efficiency (EE) of 16.36 ± 2.35% was achieved. Although lower than encapsulation efficiencies reported for other nanoparticle formulations, such as liposomal curcumin (Tefas et al., 2017; Wei et al., 2020), this value is consistent with the limited loading capacity expected for biologically derived vesicles containing endogenous cargo.

**Figure 3.**
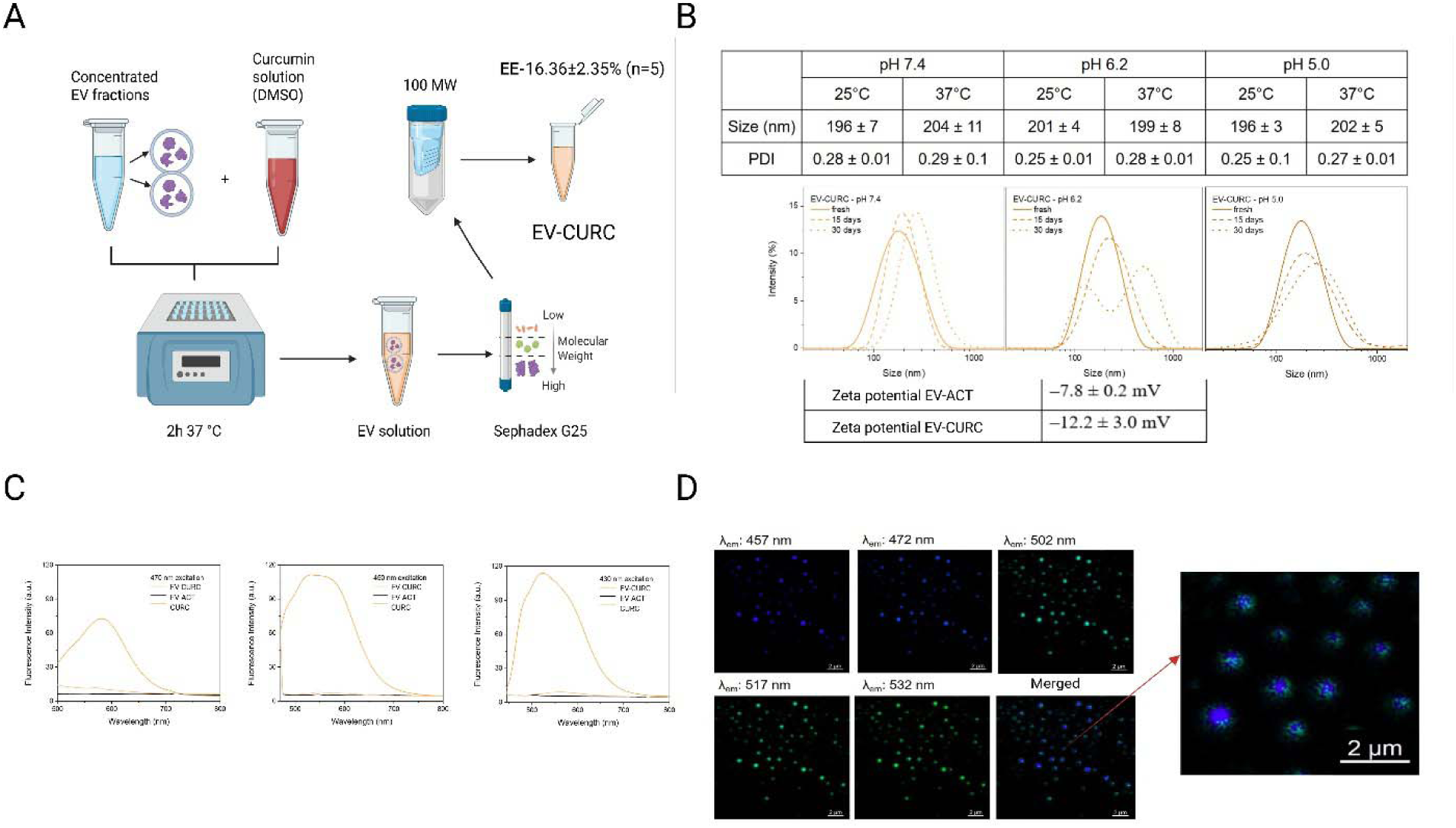
Physicochemical and fluorescence characterization of curcumin-loaded extracellular vesicles derived from activated dendritic cells (EV-CURC). **A**. Schematic representation of passive CURC loading into EV-ACT followed by size-exclusion chromatography (SEC) to separate non-associated CURC. Encapsulation efficiency (EE) was determined and calculated from five independent biological replicates (n=5), using the formula: EE (%) = (µg CURC after elution/µg CURC before elution) × 100. Data are presented as mean ± S.D. **B**. Dynamic light scattering (DLS) characterization of EV-CURC showing hydrodynamic diameter, polydispersity index (PDI), and zeta potential under different pH conditions (5.0, 6.2, and 7.4), storage times (fresh, 15 and 30 days), and temperatures (25°C and 37°C). **C**. Fluorescence emission spectra of free CURC and EV-CURC following excitation at 430, 450, and 470 nm, demonstrating distinct fluorescence profiles associated with EV-loaded CURC. **D**. Spectral confocal fluorescence microscopy of EV-CURC showing heterogeneous fluorescence emission represented as blue and green pseudocolors, consistent with spectrally distinct EV-associated CURC populations. Scale bar = 2 µm.

The hydrodynamic diameter of EV-CURC remained stable across the tested pH values (7.4, 6.2, and 5.0), with mean particle sizes ranging from 196 to 204 nm, consistent with the expected size distribution of small EVs (Welsh et al., 2024). The PDI values (0.25–0.29) further indicated a relatively homogenous particle population following CURC loading (**Figure 3B**). While measurements performed at 25°C provided information regarding storage conditions, preservation of the size distribution at 37°C demonstrates structural stability under physiological conditions (**Figure 3B**). Collectively, these findings indicate that CURC loading did not adversely affect the physicochemical stability of EVs under the tested storage and physiological conditions.

Fluorescence spectroscopy provided additional evidence of CURC association with EVs. Upon excitation at 430, 450, and 470 nm, EV-CURC displayed intense fluorescence emission spectra with broadened profiles and additional shoulders compared with free CURC (**Figure 3C**). The persistence of these spectral features across excitation wavelengths suggests the presence of at least two emissive components, indicating that EV-associated CURC resides in at least two distinct physicochemical microenvironments, potentially corresponding to membrane-associated and intravesicular curcumin populations (C. Xu et al., 2022). Zeta potential measurements further supported this observation. Empty EVs exhibited a surface charge of –7.8 ± 0.2 mV, whereas EV-CURC displayed a more negative surface potential (–12.2 ± 3.0 mV) (**Figure 3B**). This shift is consistent with partial association of CURC with the EV membrane, thereby altering the electrostatic profile of EVs (Alkhaldi et al., 2024), although intravesicular localization cannot be excluded.

Spectral confocal fluorescence microscopy further supported the presence of at least two spectrally distinct EV-associated CURC populations. Following deposition of EV-CURC onto glass slides and excitation at 405 nm, fluorescence emission was detected in multiple spectral windows represented as blue and green pseudocolors (**Figure 3D**). Taken together with the fluorescence spectroscopy and zeta potential analyses, the spectral confocal fluorescence images are consistent with the hypothesis that the blue emission predominantly originates from curcumin species associated with the EV membrane or intravesicular compartment, whereas the green emission likely reflects curcumin populations more closely associated with the external vesicle surface (**Figure 3B-D**).

Collectively, these complementary physicochemical and fluorescence analyses support the hypothesis that EV-associated curcumin exists in at least two distinct microenvironments. These observations further suggest that the spectrally distinct curcumin populations may correspond to membrane-associated and intravesicular localization, although the present data do not directly demonstrate their precise spatial distribution within EVs.

### 3.4 Curcumin-loaded EVs rapidly associate with primary CD8^+^ T cells

Efficient cellular uptake is a prerequisite for the biological activity of both free curcumin and EV-mediated cargo delivery. To evaluate the interaction of EV-associated curcumin with primary CD8**^+^** T cells in comparison with free CURC, cells were first isolated from pooled murine lymph nodes by negative selection according to the workflow shown in **Figure 4A**. Flow cytometric analysis confirmed a highly enriched CD8**^+^** T-cell population, with approximately 96% of isolated cells expressing the CD8 marker (**Figure 4B**). Following 30 min incubation of the cells with 5 µM free CURC or EV-associated CURC and removal of unbound preparations by repeated washing, fluorescence microscopy revealed a strong cell-associated CURC fluorescence signal associated with DAPI-positive CD8**^+^** T cells in both treatment groups, whereas untreated control cells exhibited only background autofluorescence (**Figure 4C**). These findings are consistent with rapid cellular association and likely uptake of both CURC and EV-associated CURC under the experimental conditions employed.

**Figure 4.**
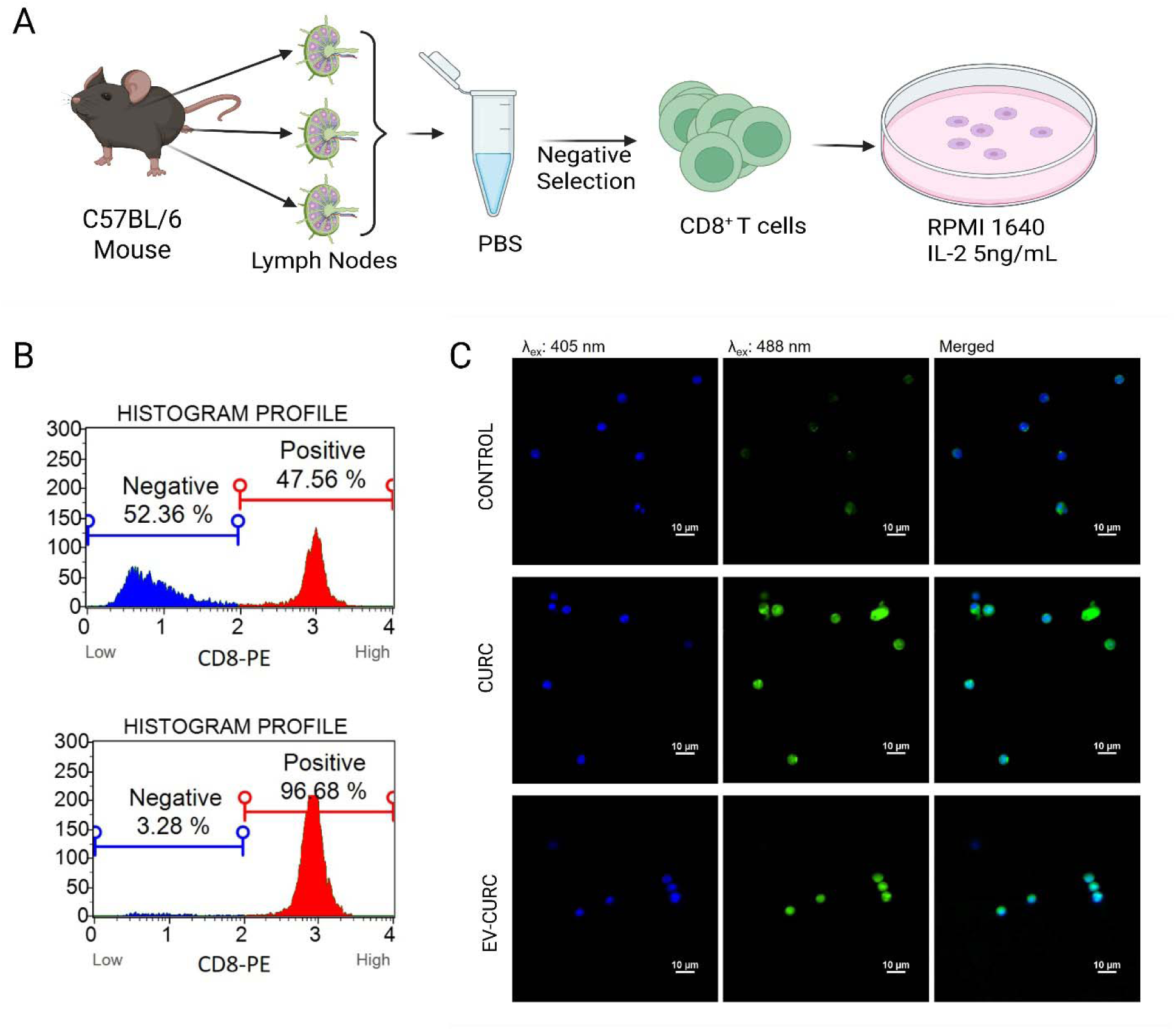
Primary CD8**^+^**T-cell purification and fluorescence microscopy analysis of EV-CURC and free CURC association with CD8**^+^** T cells. **A.** Schematic overview of CD8**^+^** T-cell isolation from pooled C57BL/6 mouse lymph nodes by negative selection. **B.** Flow cytometry analysis of CD8^+^ T cells before (upper panel) and after (lower panel) negative selection enrichment, demonstrating a CD8**^+^** T-cell purity exceeding 96%. **C.** Fluorescence microscopy of purified primary CD8**^+^** T cells following 30 min incubation with 5 µM free CURC or EV-loaded CURC (EV-CURC); CURC fluorescence is shown in green and nuclei are stained with DAPI (blue); Control = untreated CD8**^+^** T cells; CURC = CD8**^+^**T cells treated with 5 µM free curcumin; EV-CURC = CD8**^+^** T cells treated with 5 µM EV-CURC; Scale bar = 10 µm.

### 3.5 Curcumin loading selectively enhances early CD8^+^ T-cell activation while preserving EV-mediated effector-like responses

Early activation of primary CD8^+^ T cells was evaluated by flow cytometry analysis of CD69 expression following 12 and 24 h of treatment. CD69 is the earliest inducible marker of lymphocyte activation and also regulates T-cell retention and immune responses (Koyama-Nasu et al., 2022). After 12 h of treatment, treatment with EV-CURC (1 µM) significantly increased CD69 expression compared with untreated controls (*p* = 0.0083), EV-ACT (*p* = 0.0083), and free CURC (*p* = 0.0051) (**Figure 5A**). None of the remaining treatments significantly affected CD69 expression compared to untreated controls (**Figure 5A**). After 24 h of treatment, EV-CURC (1 µM) remained the only treatment that significantly increased CD69 expression compared with untreated controls (*p* = 0.0086), whereas EV-ACT and free CURC treatments failed to elicit a comparable response (**Figure 5B**). As expected, pharmacological stimulation with PMA/ionomycin, used as a receptor-independent positive control, induced the highest levels of CD69 expression at both 12 and 24 h, confirming the responsiveness of the primary CD8^+^ T cells.

**Figure 5.**
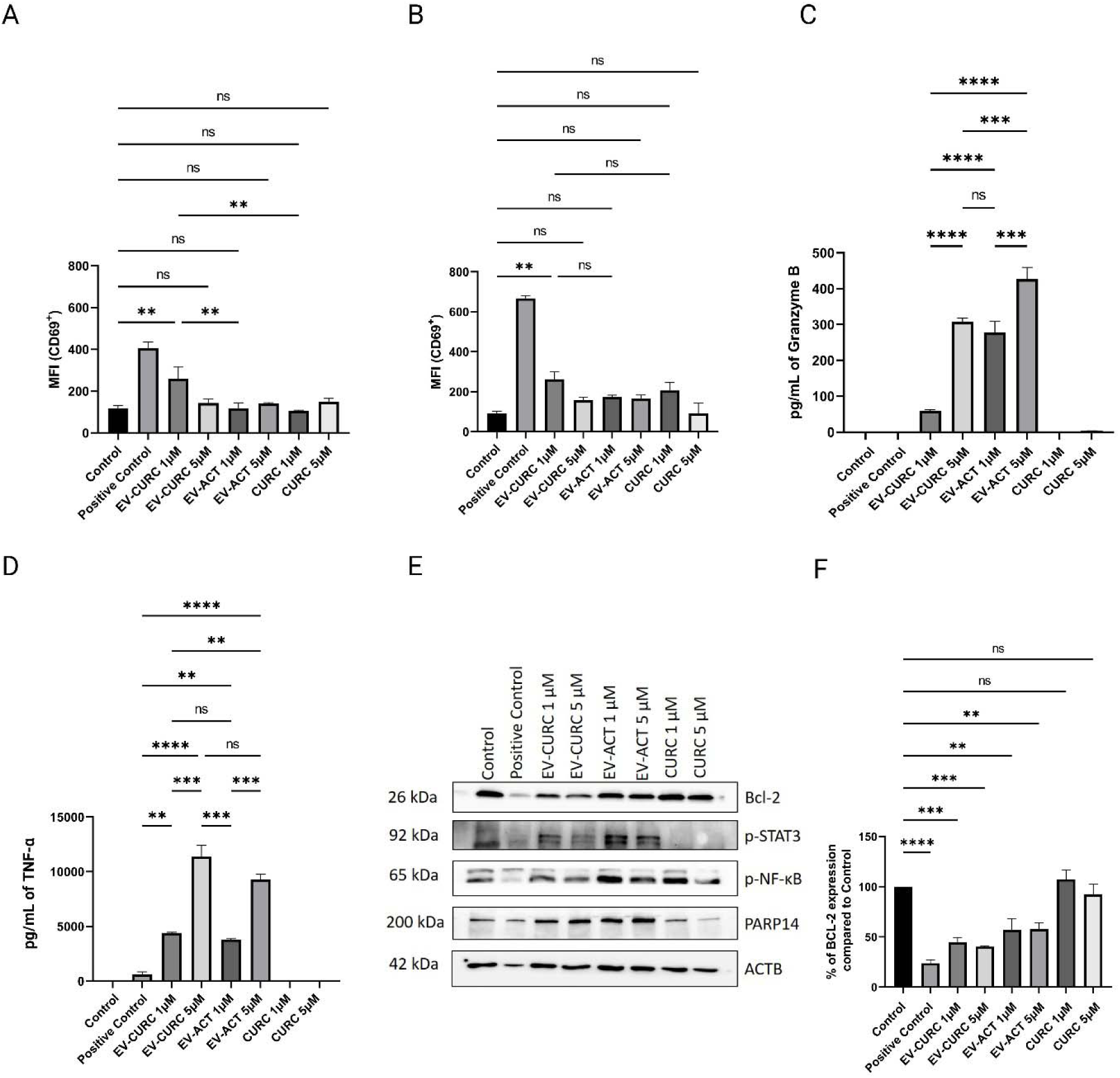
Evaluation of early activation, effector responses, and molecular changes in primary CD8^+^ T cells following treatment with EV-CURC by flow cytometry, ELISA, and western blot analysis. **A.** Flow cytometric analysis of CD69 expression in primary CD8^+^ T cells following 12 h of treatment (n = 3 technical replicates). **B.** Flow cytometric analysis of CD69 expression in primary CD8^+^ T cells following 24 h of treatment (n = 3 technical replicates). **C.** Quantification of Granzyme B secretion by ELISA in primary CD8^+^ T-cell culture supernatants following 24 h of treatment with EV-CURC, EV-ACT and free CURC (n = 2 technical replicates). **D.** Quantification of soluble TNF-α secretion by ELISA in primary CD8^+^ T-cell culture supernatants following 24 h of treatment with EV-CURC, EV-ACT and free CURC (n = 2 technical replicates). **E.** Representative western blot analysis of p-NF-κB, p-STAT3, PARP14, and Bcl-2 expression in primary CD8^+^ T cells following 24 h of treatment. β-Actin (ACTB) was used as the loading control. Densitometric quantification of p-NF-κB, p-STAT3, and PARP14 is presented in **Supplementary Figure S12A–C**. **F.** Densitometric quantification of relative Bcl-2 protein expression normalized to ACTB following 24 h of treatment (n = 2 technical replicates). Statistical significance was determined by one-way ANOVA followed by Dunnett’s post hoc test. Data are presented as mean ± S.D.; ns – not significant; P > .05; *, P < .05; **, P< .01; ***, P <.001; ****, P <.0001.

Because CD69 expression was reported to peak around 24 h following T-cell activation (Ziegler et al., 1994), coinciding with early acquisition of cytotoxic effector functions including the production of canonical effector molecules such as Granzyme B, TNF-α, and IFN-γ (Mardiney et al., 1996; Zimmermann et al., 1999), we subsequently evaluated the secretion of Granzyme B and TNF-α in primary CD8^+^ T-cell culture supernatants following 24 h of treatment using ELISA. Compared with untreated controls, activated dendritic cell-derived EVs significantly promoted Granzyme B secretion, whereas free CURC at either concentration failed to induce detectable changes. Among the EV-CURC-treated groups, only the 5 µM concentration significantly increased Granzyme B secretion (**Figure 5C**). Specifically, Granzyme B secretion was significantly increased following CD8^+^ T-cell treatment with EV-CURC 5 µM (*p* < 0.0001), EV-ACT 1 µM (*p* < 0.0001), and EV-ACT 5 µM (*p* < 0.0001) compared with untreated controls. Both EV formulations exhibited concentration-dependent increases in Granzyme B secretion, with significantly higher levels detected at 5 µM than at the corresponding 1 µM concentration. Notably, EV-ACT elicited significantly stronger Granzyme B secretion than EV-CURC at the corresponding concentrations (**Figure 5C**). A similar pattern was observed for TNF-α secretion when CD8^+^ T cells were treated with both EV-CURC (1 and 5 µM) and EV-ACT (1 and 5 µM) as TNF-α production was detectable, whereas untreated cells and cells treated with free CURC exhibited cytokine concentrations below the assay detection limit (**Figure 5D**). Compared with the positive control, TNF-α secretion remained significantly lower following treatment with EV-CURC (1 µM, *p* = 0.0040; 5 µM, *p* < 0.0001) and EV-ACT (1 µM, *p* = 0.0087; 5 µM, *p* < 0.0001). No statistically significant differences were observed between EV-CURC and EV-ACT at the corresponding concentrations (**Figure 5D**).

Collectively, these findings demonstrate that EV-ACT promote early activation and acquisition of effector functions in primary CD8^+^ T cells. Whereas curcumin loading selectively enhanced early CD69 expression on CD8^+^ T cells, activated EVs alone induced the strongest Granzyme B release, while both EV formulations comparably stimulated TNF-α secretion with slightly higher levels observed in the cells receiving EV-CURC.

### 3.6 EV-CURC selectively modulates early signaling pathways associated with CD8^+^ T-cell activation while preserving intrinsic EV-mediated effects

To gain mechanistic insight into the signaling events underlying EV-mediated early activation of primary CD8^+^ T cells, we investigated the expression of proteins with established roles in regulating T-cell activation, effector differentiation, cytokine production, and cell survival by western blot analysis. Specifically, we assessed proteins involved in inflammatory signaling (p-NF-κB) (Daniels & Teixeiro, 2025), T-cell activation and differentiation (p-STAT3) (Imataki et al., 2012), T-cell transcriptional regulation (PARP14) (Mehrotra et al., 2011), and cell survival remodeling (Bcl-2) (Bhattacharyya et al., 2007), following 24 h of treatment (**Figure 5E and F**, **Supplementary Figures S12A-C**).

Among the signaling proteins evaluated, STAT3 phosphorylation showed the most pronounced response to EV-mediated curcumin delivery. EV-CURC significantly increased p-STAT3 at both concentrations tested compared with untreated controls (EV-CURC 1 µM, *p* = 0.0471; EV-CURC 5 µM, *p* = 0.0141), whereas free CURC had no significant effect (**Figure 5E and Supplementary Figure S12B**). These findings indicate that EV-mediated delivery selectively promotes STAT3 activation during early CD8^+^ T-cell activation. In contrast, p-NF-κB was only modestly but not significantly affected by the EV treatments. A significant increase was detected only following treatment with free CURC 1 µM (*p* = 0.0202) compared to control which is consistent with previous studies (Zheng et al., 2025b), whereas p-NF-κB levels remained unchanged in the EV-treated groups (**Figure 5E and Supplementary Figure S12A**). PARP14 expression was increased following EV-based treatments compared to control, with the highest levels observed in EV-CURC-treated cells, followed by EV-ACT, whereas free CURC-treated cells displayed lower levels compared to control (**Figure 5E and Supplementary Figure S12C**). Both EV-CURC and EV-ACT significantly reduced Bcl-2 expression compared with untreated controls (EV-CURC 1 µM, *p* = 0.0003; EV-CURC 5 µM, *p* = 0.0002; EV-ACT 1 µM, *p* = 0.0015; EV-ACT 5 µM, *p* = 0.0017), whereas free CURC had no significant effect (**Figure 5E and F**). Despite the reduction in Bcl-2 expression in EV-treated conditions, no corresponding decrease in cell viability was observed (**Supplementary Figure S12D**), indicating that the reduced Bcl-2 levels were not associated with overt cell death under the experimental conditions employed.

Collectively, these findings demonstrate that EV-mediated curcumin delivery selectively modulates early signaling pathways associated with CD8^+^ T-cell activation. Compared with EV-ACT alone, EV-CURC preferentially enhanced STAT3 phosphorylation and exhibited the highest PARP14 expression while reducing Bcl-2 levels without compromising cell viability, supporting a distinct early molecular response to EV-delivered curcumin.

## Discussion

Extracellular vesicles (EVs) are versatile nanoscale drug delivery systems, owing to their intrinsic biocompatibility, ability to incorporate bioactive cargo, and natural capacity to interact with recipient cells (Dang et al., 2020). Beyond their potential as delivery vehicles, EVs derived from dendritic cells are increasingly recognized as cell-free antigen-presenting platforms because they retain immune-associated molecules inherited from their parental cells (Kowal & Tkach, 2019; Moya-Guzmán et al., 2023; Pitt et al., 2016). However, whether engineering activated dendritic cell-derived EVs with immunomodulatory cargo can further influence the early molecular and functional events governing CD8^+^ T-cell activation has remained largely unexplored. In the present study, we demonstrated that EVs derived from CpG-activated and TRP-2 peptide-pulsed dendritic cells possess an intrinsic immunostimulatory phenotype that can be selectively potentiated through curcumin loading, resulting in enhanced early activation accompanied by modulation of key signaling proteins in primary CD8^+^ T cells.

The EV isolation strategy employed herein combined ultrafiltration with size-exclusion chromatography (UF-SEC) (**Figure 1B**), a robust approach for enriching small EVs while minimizing soluble protein contamination, particle aggregation while preserving vesicle integrity (Benedikter et al., 2017; Ter-Ovanesyan et al., 2021). Consistent with current MISEV2023 recommendations (Théry et al., 2018; Welsh et al., 2024), the isolated vesicles exhibited physicochemical properties characteristic of small EVs (∼170 nm) and expressed canonical EV-associated proteins while lacking the Golgi marker GM130 (**Figure 1C-E**).

Importantly, exploratory proteomic profiling further revealed that EVs released from activated dendritic cells possessed a distinct immune-associated protein repertoire characterized by coordinated enrichment in molecules involved in antigen processing (CALR and CSTA) (Kasikova et al., 2019; Nasti et al., 2025), MHC-I-associated pathways (Tian et al., 2007), endoplasmic reticulum protein folding, immunoproteasome function, vesicle trafficking, and immune adhesion (ITGA4, ICAM1 and ITGB1) (Bergqvist et al., 2025; Makgoba et al., 1988) (**Figure 2C and D**). Although these findings do not directly demonstrate antigen presentation by EV-ACT, they suggest that activation remodels several complementary steps of the antigen presentation machinery rather than individual proteins (Pérez-Montesinos et al., 2017). These observations are consistent with previous reports indicating that dendritic cell-derived EVs retain immune-associated molecular components capable of supporting immune communication and cross-presentation-related processes (Curtsinger et al., 1999; Wahlund et al., 2017) as biologically active immunomodulatory particles rather than passive delivery vehicles (Théry et al., 2002; Tkach et al., 2017).

Curcumin was selected as the exogenous cargo because of its well-documented immunomodulatory and anti-tumor properties at low doses (L. Liu et al., 2021b) (Limsakul et al., 2025; Zeynali et al., 2025) together with its limited therapeutic potential resulting from poor aqueous solubility and low systemic bioavailability (Wei et al., 2020; Zheng et al., 2025b). Passive curcumin loading resulted in an encapsulation efficiency of approximately 16%, which is expected for biologically derived vesicles already containing endogenous cargo (Théry et al., 2001). Despite this modest loading efficiency, EV-CURC (∼200 nm) shifted toward increased electronegative zeta potential, change consistent with successful cargo association (M. S. Kim et al., 2016), retained physicochemical stability under both storage and physiological conditions (**Figure 3**). Fluorescence spectroscopy, zeta potential measurements, and spectral confocal microscopy suggested that EV-associated curcumin occupies at least two physicochemical microenvironments (**Figure 3**). Although the precise localization of curcumin within EVs was not determined, previous studies have suggested that both membrane-associated and intravesicular distribution were possible (Shie et al., 2025).

Both free curcumin and EV-delivered curcumin rapidly associated with primary CD8^+^ T cells within 30 min (**Figure 4**), consistent with previous reports describing efficient cellular interaction of EV-delivered curcumin (C. Xu et al., 2022). Although the imaging approach used here does not directly demonstrate intracellular localization, these observations indicate efficient cell association under the experimental conditions employed and are compatible with previously described mechanisms of EV uptake (Costa Verdera et al., 2017).

One of the principal findings of the present study is that curcumin loading did not uniformly amplify all biological responses induced by activated dendritic cell-derived EVs, but rather selectively modulated specific components of the early activation program. Evaluation of the early activation marker CD69 in CD8^+^ T cells (Simms & Ellis, 1996) demonstrated that EV-CURC selectively increased and sustained CD69 expression, whereas activated EVs alone and free curcumin failed to elicit a comparable response (**Figure 5A and B**). This suggests that EV-mediated curcumin delivery exerts immunomodulatory effects by preferentially reinforcing the earliest activation events rather than globally increasing T-cell stimulation (Sun et al., 2010).

Interestingly, this selective modulation extended to downstream functional responses. Both EV-ACT and EV-CURC promoted Granzyme B and TNF-α secretion, whereas free curcumin alone remained largely ineffective. However, activated EVs consistently induced stronger Granzyme B production than EV-CURC at equivalent concentrations, while TNF-α secretion remained comparable between the two EV preparations with a tendency toward slightly higher levels following EV-CURC treatment (**Figure 5C and D**). Rather than indicating reduced biological activity of EV-CURC, these findings suggest that engineering EV cargo does not necessarily potentiate every downstream effector function. Instead, curcumin appears to selectively reshape specific aspects of the early activation program while preserving the intrinsic immunostimulatory capacity of activated dendritic cell-derived EVs.

The molecular analyses further support this concept by revealing selective rather than global modulation of pathways associated with effector-like cell differentiation rather than broad inflammatory activation. Among the signaling proteins evaluated, STAT3 phosphorylation exhibited the most pronounced response to EV-mediated curcumin delivery, whereas free curcumin produced no detectable effect (**Supplementary Figure S12**). Although STAT3 exerts context-dependent functions during T-cell differentiation (Cui et al., 2011; Imataki et al., 2012; Kane et al., 2014), the increased STAT3 phosphorylation observed here occurred together with enhanced CD69 expression and preserved production of TNF-α and Granzyme B. Within the early activation context investigated here, the data are consistent with STAT3 activation as part of an early activation-associated signaling response rather than an immunosuppressive phenotype. Similarly, the expression of PARP14, an important mediator of immune-cell activation and inflammatory responses (Mehrotra et al., 2011), was consistently highest following EV-CURC treatment (**Supplementary Figure S12**). Given that PARP14 has been described as a transcriptional co-regulator of STAT-dependent gene expression (Mehrotra et al., 2011, 2015), these observations suggest coordinated modulation of signaling events associated with early T-cell activation, although causal relationships between these molecular changes and the functional phenotype remain to be established.

By comparison, NF-κB phosphorylation was only modestly affected by EV-based treatments. Because canonical NF-κB activation is highly transient, peaking within the first hours following T-cell stimulation before declining substantially (Auphan et al., 1999), the modest changes observed at 24 h were not unexpected, despite the central role of NF-κB in regulating T-cell activation, inflammatory responses, and survival (Gerondakis et al., 2014). Likewise, the reduction in Bcl-2 expression observed following EV treatments occurred without any measurable loss of cell viability (**Figure 5E and F**, **Supplementary Figure S12**), suggesting that the observed Bcl-2 downregulation reflects physiological remodeling of survival pathways accompanying the transition of CD8^+^ T cells toward an effector phenotype, rather than apoptosis induction (Boise et al., 1995; Hildeman et al., 2007).

The present work focused on the earliest molecular and functional events elicited by activated dendritic cell-derived EVs, thereby providing a framework for understanding the initial stages of CD8^+^ T-cell activation that precede differentiation. Whether these early molecular changes ultimately translate into durable cytotoxic responses and improved anti-tumor immunity remains to be established. Nevertheless, the present findings provide proof-of-concept that engineering biologically active dendritic cell-derived EVs with immunomodulatory cargo can selectively reprogram the earliest stages of CD8^+^ T-cell activation while preserving the intrinsic immunostimulatory properties of the parental vesicles.

## Conclusions

Collectively, our findings support the use of activated dendritic cell-derived EVs as intrinsically immunostimulatory platforms that can be selectively enhanced by immunomodulatory cargo loading. Their endogenous immune-associated proteomic cargo promotes early activation and effector responses in primary CD8^+^ T cells, whereas curcumin selectively reinforces early activation-associated molecular programs without compromising cell viability. Together, these findings provide proof-of-concept that engineering biologically active dendritic cell-derived EVs with immunomodulatory cargo can selectively fine-tune early CD8^+^ T-cell responses while preserving the intrinsic immunostimulatory properties of the parental vesicles, thereby supporting the further development of EV-based immunotherapies.

## Author Contributions

**S.M.D.**: Methodology, Investigation, Data Curation, Formal Analysis, Validation, Writing Original Draft, **L.P.**: Investigation, Validation, Data Curation, Writing, Review & Editing, **S.M.**: Methodology, Validation, Data Curation, **C.V.A.M.**: Investigation, Methodology, Formal Analysis, Resources, **O.I.P.**: Methodology, **R.B.**: Investigation, Methodology, **M.F.**: Investigation, Methodology, **A.B.M.**: Methodology, **A.M.**: Methodology, **L.S.**: Investigation,, Methodology, **M.B.**: Conceptualization, Supervision, Writing, Review & Editing Supervision, **A.S.**: Conceptualization, Supervision, Writing, Review & Editing, Funding Acquisition, Project Administration.

## Data Availability Statement

The mass spectrometry proteomics data have been deposited to the MassIVE repository (University of California San Diego) under accession number MSV000101874 (MassIVE Private Dataset). The dataset is currently private and will be made available after publication.

The data supporting the findings of this study are available from the corresponding author upon reasonable request.

## Declaration of generative AI use

The authors acknowledge the use of generative AI (ChatGPT 3.5) to improve grammar, language and readability of the manuscript. All text was critically reviewed and edited by the authors, who take full responsibility for the final manuscript.

