## Supplementary Table S2 for "Extracellular Vesicles Derived from Activated Dendritic Cells Loaded with Curcumin Promote Early Activation-associated Functional and Molecular Reprogramming of Primary CD8^+^ T Cells"

**Table 2.** Mass spectrometry identified proteins in EV-NA and EV-ACT that were not previously reported in ExoCarta and EVpedia for murine proteins.

| **Gene Symbol EV-NA** | **Gene Symbol EV-ACT** |
| --- | --- |
| H4C1 | H4C1 |
| H2BC3 | H2BC3 |
| H2AC25 | H2AC25 |
| HARS1 | HARS1 |
| GBA1 | GBA1 |
| SEPTIN2 | SEPTIN2 |
| RARS1 | RARS1 |
| KARS1 | KARS1 |
| H2AZ2 | H2AZ2 |
| CYRIB | CYRIB |
| SEPTIN7 | SEPTIN7 |
| GFUS | GFUS |
| H3-3A; H3-3B | H3-3A; H3-3B |
| VARS1 | VARS1 |
| SEPTIN11 | SEPTIN11 |
| GARS1 | GARS1 |
| TARS1 | TARS1 |
| ADPRS | ADPRS |
| ATP5F1A | ATP5F1A |
| EPRS1 | EPRS1 |
| FBXL21 | FBXL21 |
| PTS | PTS |
| TIMM8A2 | TIMM8A2 |
| NDP | NDP |
| YARS1 | YARS1 |
| ATP5F1B | ATP5F1B |
| PDXP | PDXP |
| ZNG1 | ZNG1 |
| ASRGL1 | ASRGL1 |
| ENOPH1 | ENOPH1 |
| SKIC8 | SKIC8 |
| CARS1 | CARS1 |
| PRMT6 | PRMT6 |
| UXT | UXT |
| DARS1 | DARS1 |
| POLR2G | POLR2G |
| S100A14 | S100A14 |
| GGCT | GGCT |
| CTBS | CTBS |
| APIP | APIP |
| VIL1 | VIL1 |
| LRFN5 | LRFN5 |
| INSRR | INSRR |
| RNASET2B | RNASET2B |
| TTC21A | TTC21A |
| CHD6 | CHD6 |
| PLPBP | PLPBP |
| GPT | GPT |
| BEND3 | BEND3 |
| MIS18BP1 | MIS18BP1 |
| IARS1 | IARS1 |
| AOC2 | AOC2 |
| SBNO2 | SBNO2 |
| NQO2 | NQO2 |
| SLFN2 | SLFN2 |
| FITM1 | FITM1 |
| PHB1 | PHB1 |
| CA3 | CA3 |
| FAM81A | FAM81A |
| NARS1 | NARS1 |
| TTC33 | TTC33 |
| PYCR3 | PYCR3 |
| DNAAF10 | DNAAF10 |
| DCTPP1 | DCTPP1 |
| GET3 | GET3 |
| REV1 | REV1 |
| TRMT10A | TRMT10A |
| ELFN2 | ELFN2 |
| SEMA3G | SEMA3G |
| AARD | AARD |
| NFKBIL1 | NFKBIL1 |
| TDRD6 | TDRD6 |
| TBC1D30 | TBC1D30 |
| PCDH12 | PCDH12 |
| FCRL5 | FCRL5 |
| RAD52 | RAD52 |
| CYP2U1 | CYP2U1 |
| MMRN2 | MMRN2 |
| XXYLT1 | XXYLT1 |
| KRT26 | KRT26 |
| EML5 | EML5 |
| KRT12 | KRT12 |
| CDH6 | CDH6 |
| CNKSR2 | CNKSR2 |
| IGDCC4 | IGDCC4 |
| ATP5MF | ATP5MF |
| DIABLO | DIABLO |
| ADGRV1 | ADGRV1 |
| ADCK5 | ADCK5 |
| TOGARAM1 | TOGARAM1 |
| ELL3 | ELL3 |
| DNASE1L2 | LARS1 |
| HIF3A | CYCT |
| ADISSP | GBP6 |
| EVA1C | CRYL1 |
| HELQ | IWS1 |
| MIEN1 | KRT80 |
| CCDC40 | DTD1 |
| RBL2 | HMGN5 |
| KNL1 | CELSR3 |
| LMAN1L | B3GAT1 |
| SIK2 | CYP11A1 |
|  | CCK |
|  | E2F7 |
|  | SLC22A16 |
|  | SLC51A |
|  | RIPOR3 |
|  | INHCA |
|  | AMOTL1 |
|  | EFHD1 |
|  | CEP135 |
|  | KLHDC9 |
|  | GSDMA2 |
|  | BTBD7 |
|  | MRTFA |
|  | CCDC65 |
|  | FECH |
|  | KDM6A |
|  | HROB |
