## Supplementary Table S1 for "Extracellular Vesicles Derived from Activated Dendritic Cells Loaded with Curcumin Promote Early Activation-associated Functional and Molecular Reprogramming of Primary CD8^+^ T Cells"

**Supplementary Table S1.** Information on the antibodies used for western blot and flow cytometry analyses.

| **Target protein** | **Antibody type** | **Host species** | **Dilution** | **Application** | **Supplier** | **Catalog number** |
| --- | --- | --- | --- | --- | --- | --- |
| TSG101 | Primary | Mouse mAb | 1:1000 | WB | Thermo Fisher Scientific | MA5-32463 |
| STOM | Primary | Mouse mAb | 1:1000 | WB | Santa Cruz Biotechnology | sc-376869 |
| Gal3BP | Primary | Mouse mAb | 1:1000 | WB | Santa Cruz Biotechnology | sc-374541 |
| HSP70 | Primary | Mouse mAb | 1:2500 | WB | Novus Biologicals | NB-120-2788 |
| FASN | Primary | Mouse mAb | 1:1000 | WB | Santa Cruz Biotechnology | sc-48357 |
| GM130 | Primary | Mouse mAb | 1:1000 | WB | Santa Cruz Biotechnology | sc-55591 |
| Bcl-2 | Primary | Mouse mAb | 1:1000 | WB | Santa Cruz Biotechnology | Sc-7382 |
| p-NF-κB | Primary | Mouse mAb | 1:1000 | WB | Santa Cruz Biotechnology | Sc-166748 |
| PARP-14 | Primary | Mouse mAb | 1:1000 | WB | Santa Cruz Biotechnology | Sc-377150 |
| p-STAT3 | Primary | Mouse mAb | 1:1000 | IHC-P/IF | Bioss Inc. | BSM-33301M |
| Horseradish peroxidase (HRP)-conjugated IgG | Secondary | Goat α-Rabbit | 1:2500 | WB | Santa Cruz Biotechnology | sc-2004 |
| Horseradish peroxidase (HRP)-conjugated IgG | Secondary | Goat α-Mouse | 1:2500 | WB | Santa Cruz Biotechnology | sc-2005 |
| CD86 | PE-labeled | Rat IgG2B | 1:200 | IF | BioLegend | 105008 |
| MHC class II | PE-Cy5-labeled | Rat IgG2B | 1:200 | IF | BioLegend | 107612 |
| TRP-2 | Primary | Mouse mAb | 1:200 | WB/IF | Santa Cruz Biotechnology | sc-74439 |
| MHC class I | Primary | Rabbit  pAb | 1:200 | WB/IF | Thermo Fisher Scientific | MA5-48095 |
| PE-conjugated | Secondary | Goat α-Mouse | 1:300 | WB/IF | Invitrogen | A11002 |
| PE-Cy5-conjugated | Secondary | Goat α-Rabbit | 1:300 | WB/IF | Invitrogen | L42018 |
