## Supplementary Information for "Extracellular Vesicles Derived from Activated Dendritic Cells Loaded with Curcumin Promote Early Activation-associated Functional and Molecular Reprogramming of Primary CD8^+^ T Cells"

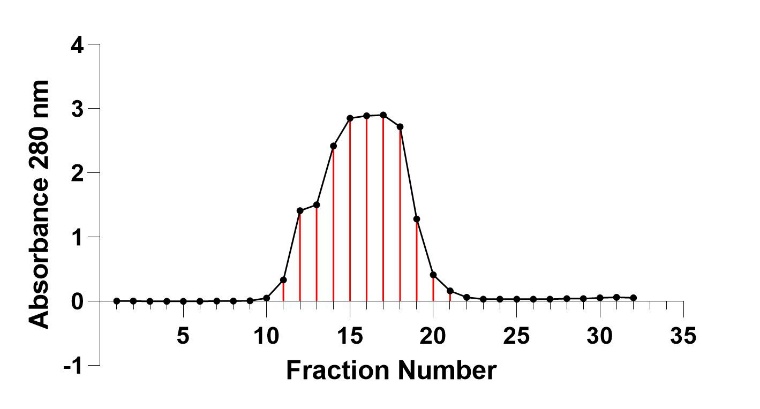

**Supplementary Figure S1.** Elution of EVs following size-exclusion chromatography using the Sephacryl S-200 HR column and representative fractions collected for further analyses. Fractions collected from the SEC collection and validation of EV protein at 280 nm. A total of 32 fractions were eluted and their absorbance was monitored at 280 nm. The EV representative fractions (11-21) were selected for further analysis.

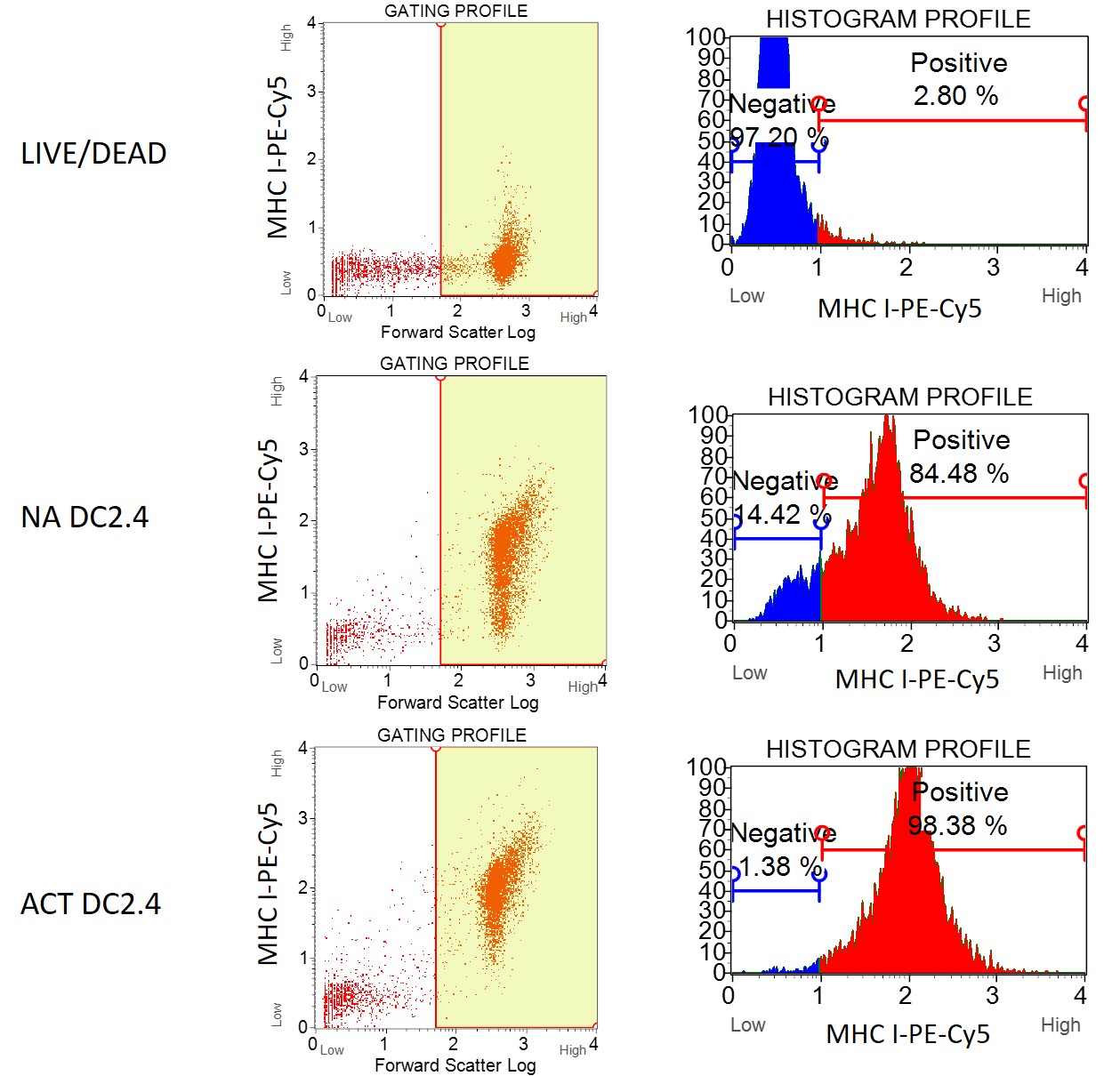

**Supplementary Figure S2.** Representative flow cytometry plots showing viability staining and MHC-I expression in non-activated (NA) DC2.4 cells and DC2.4 cells following CpG activation and TRP-2 peptide-pulsing (ACT). The plots show gating strategy with Forward Scatter Log (left) gating profile and histogram profiles (right) after staining with anti-MHC-I-PE-Cy5 antibody.

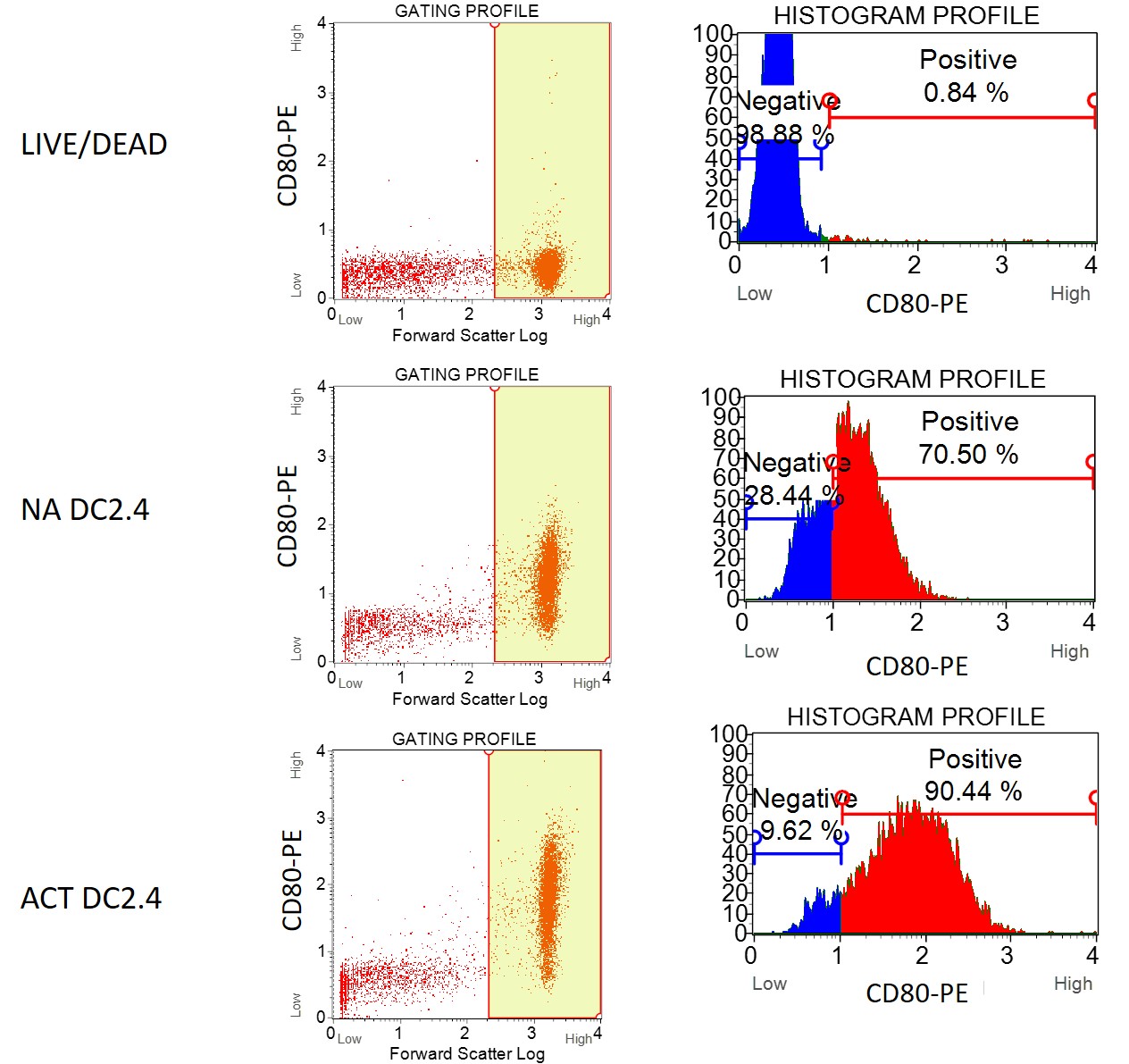

**Supplementary Figure S3.** Representative flow cytometry plots showing viability staining and CD80 expression in non-activated (NA) DC2.4 cells and DC2.4 cells following CpG activation and TRP-2 peptide-pulsing (ACT). The plots show gating strategy with Forward Scatter Log (left) gating profile and histogram profiles (right) after staining with anti-CD80-PE antibody.

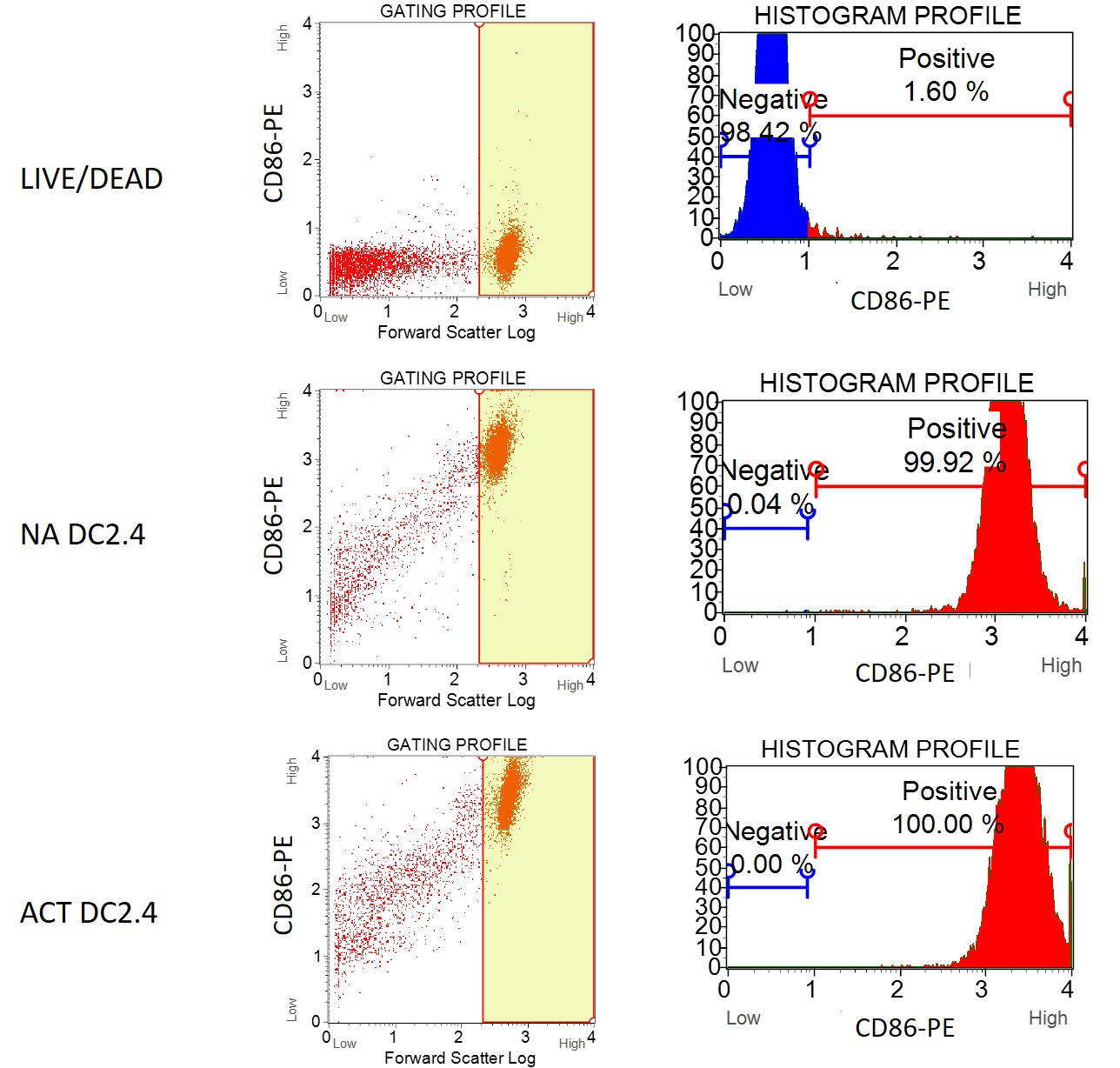

**Supplementary Figure S4.** Representative flow cytometry plots showing viability staining and CD86 expression in non-activated (NA) DC2.4 cells and DC2.4 cells following CpG activation and TRP-2 peptide-pulsing (ACT). The plots show gating strategy with Forward Scatter Log (left) gating profile, percentages of positive cells (middle) and histogram profiles (right) after staining with anti-CD86-PE antibody.

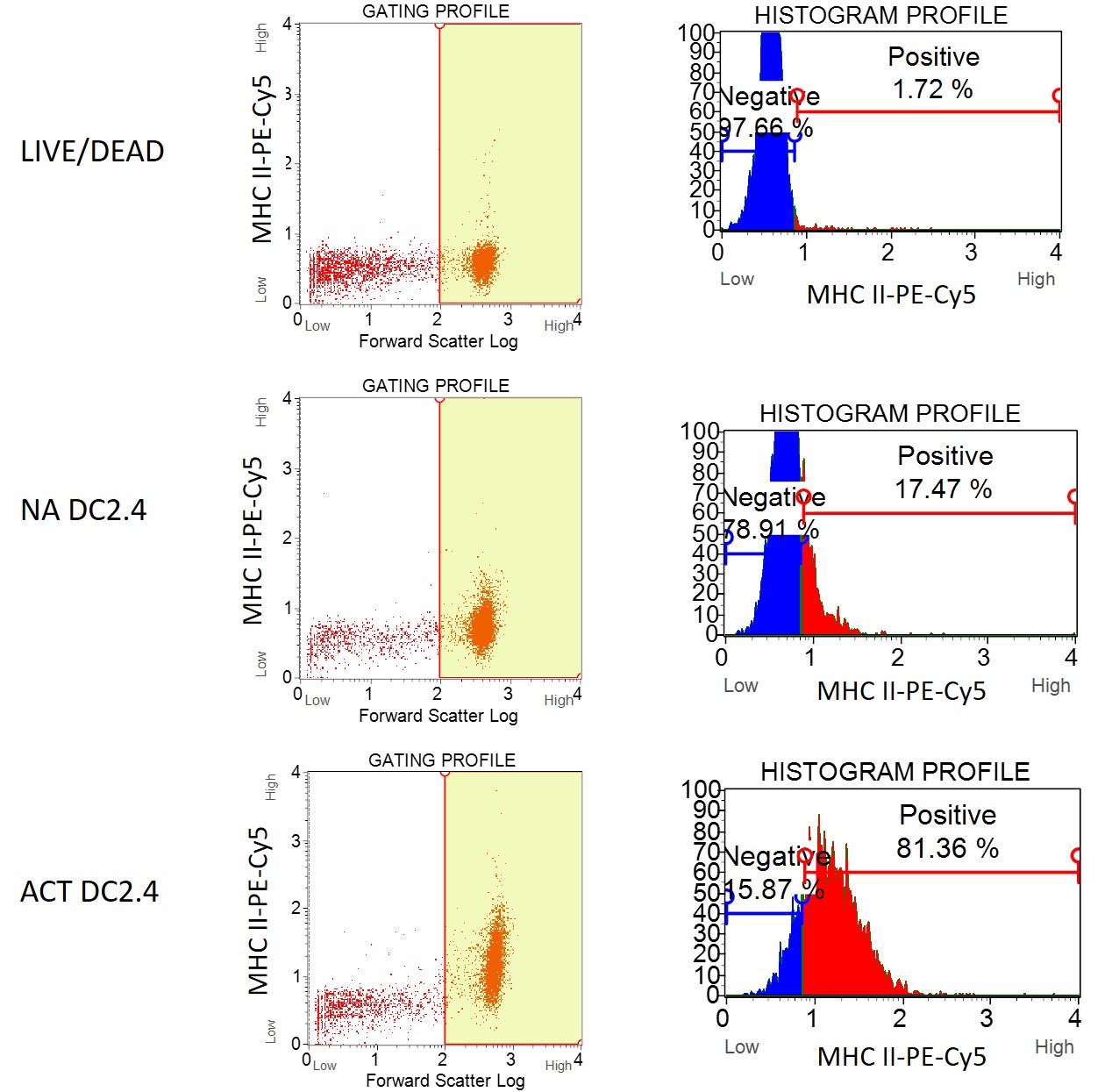

**Supplementary Figure S5.** Representative flow cytometry plots showing viability staining and MHC-II expression in non-activated (NA) DC2.4 cells and DC2.4 cells following CpG activation and TRP-2 peptide-pulsing (ACT). The plots show gating strategy with Forward Scatter Log (left) gating profile and histogram profiles (right) after staining with anti-MHC-II-PE-Cy5 antibody.

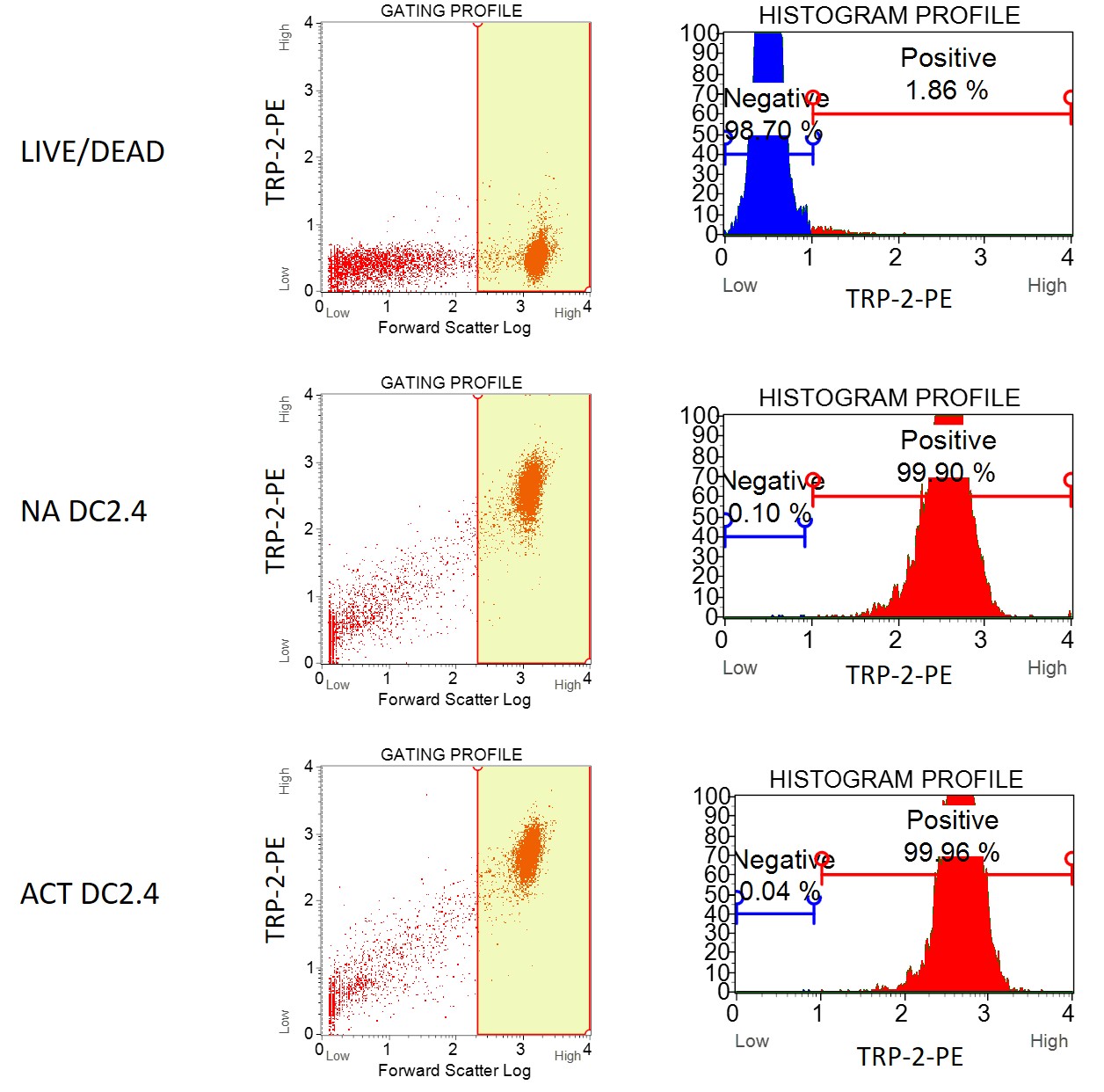

**Supplementary Figure S6.** Representative flow cytometry plots showing viability staining and TRP-2 expression in non-activated (NA) DC2.4 cells and DC2.4 cells following CpG activation and TRP-2 peptide-pulsing (ACT). The plots show gating strategy with Forward Scatter Log (left) gating profile and histogram profiles (right) after staining with anti-TRP-2-PE antibody.

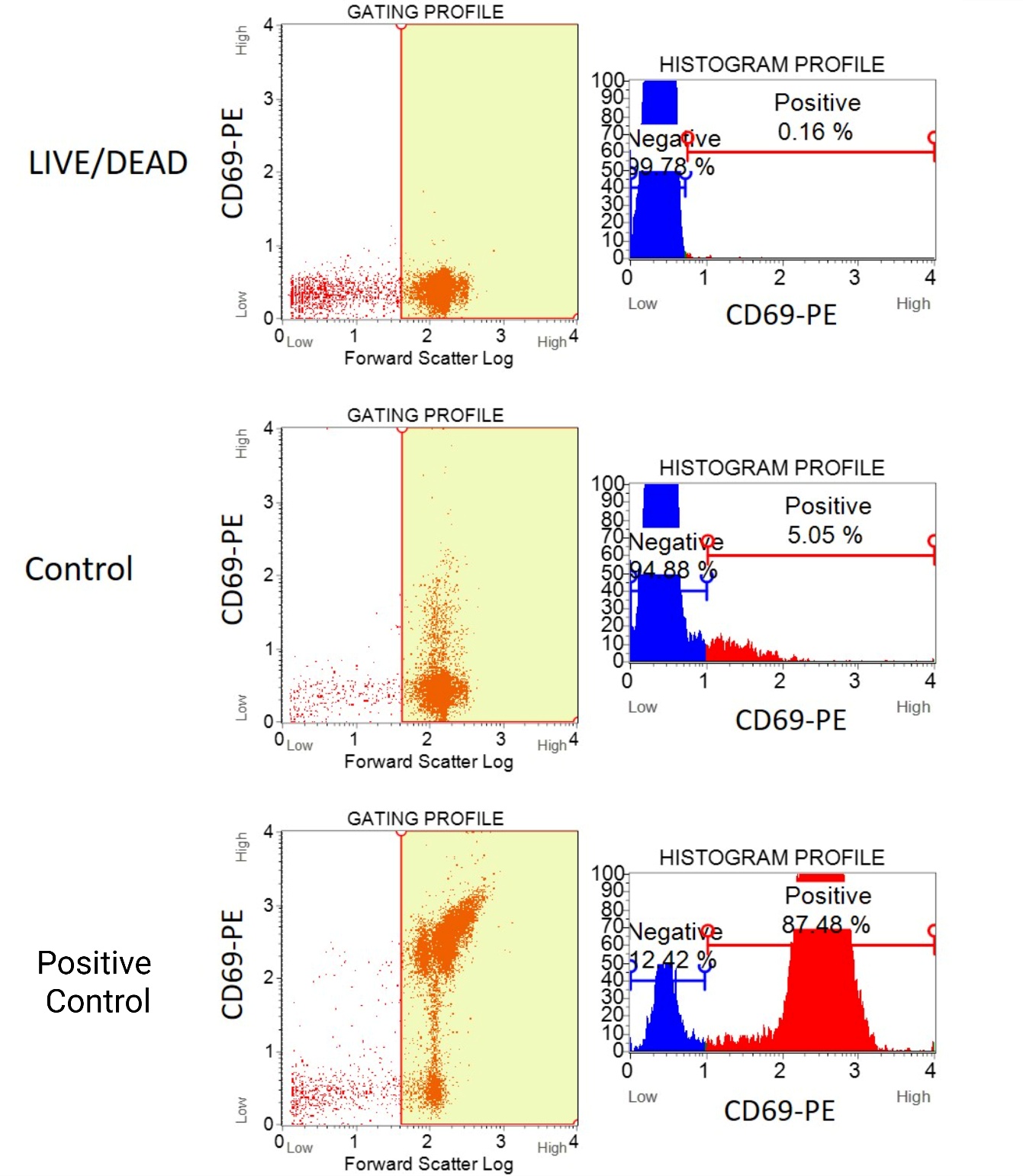

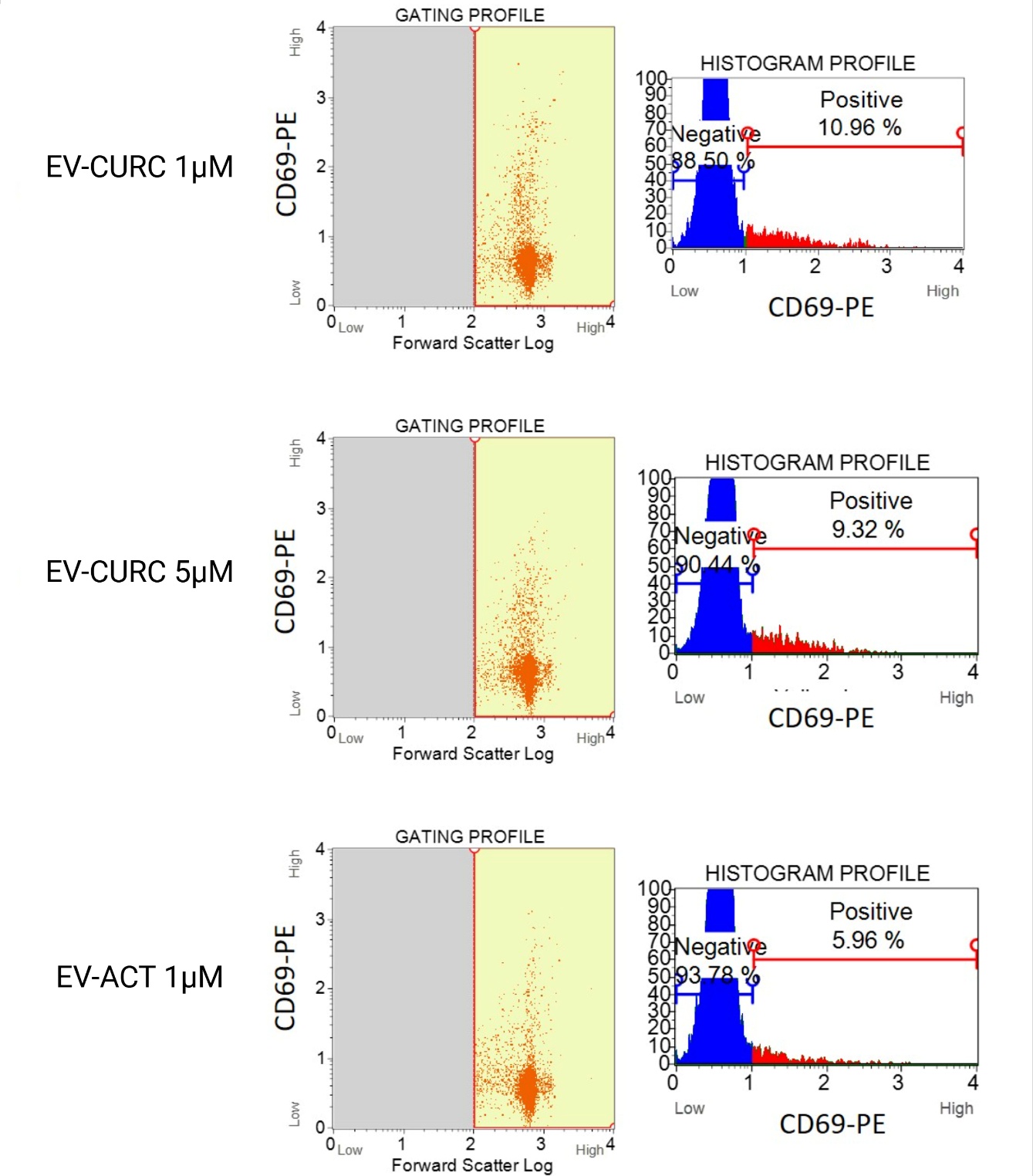

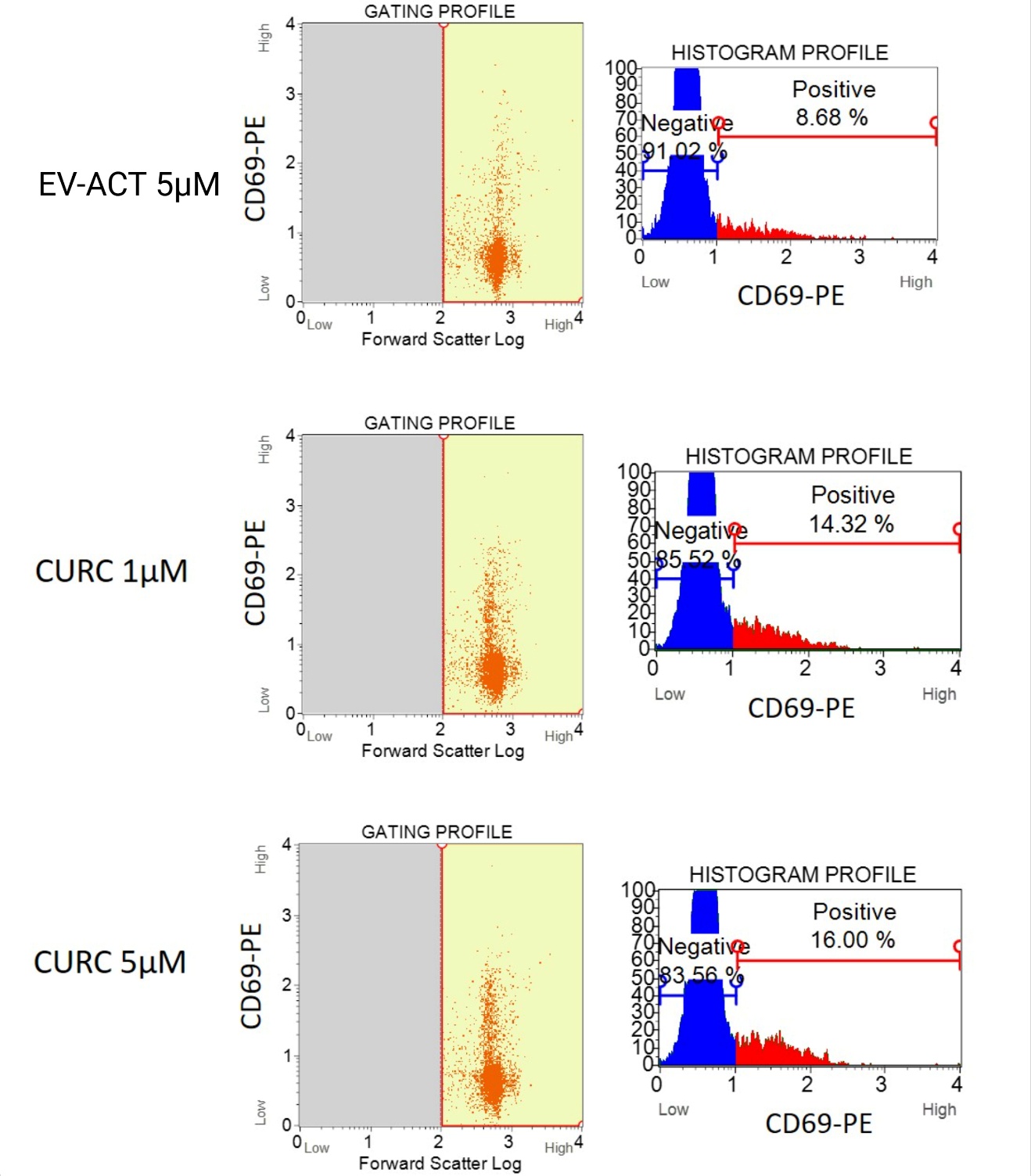

**Supplementary Figure S7.** Representative flow cytometry plots showing viability staining and CD69 expression in CD8^+^ T cells after 12 h treatment with EV-CURC. The plots show gating strategy after staining with anti-CD69-PE (Left) and corresponding histograms (Right).

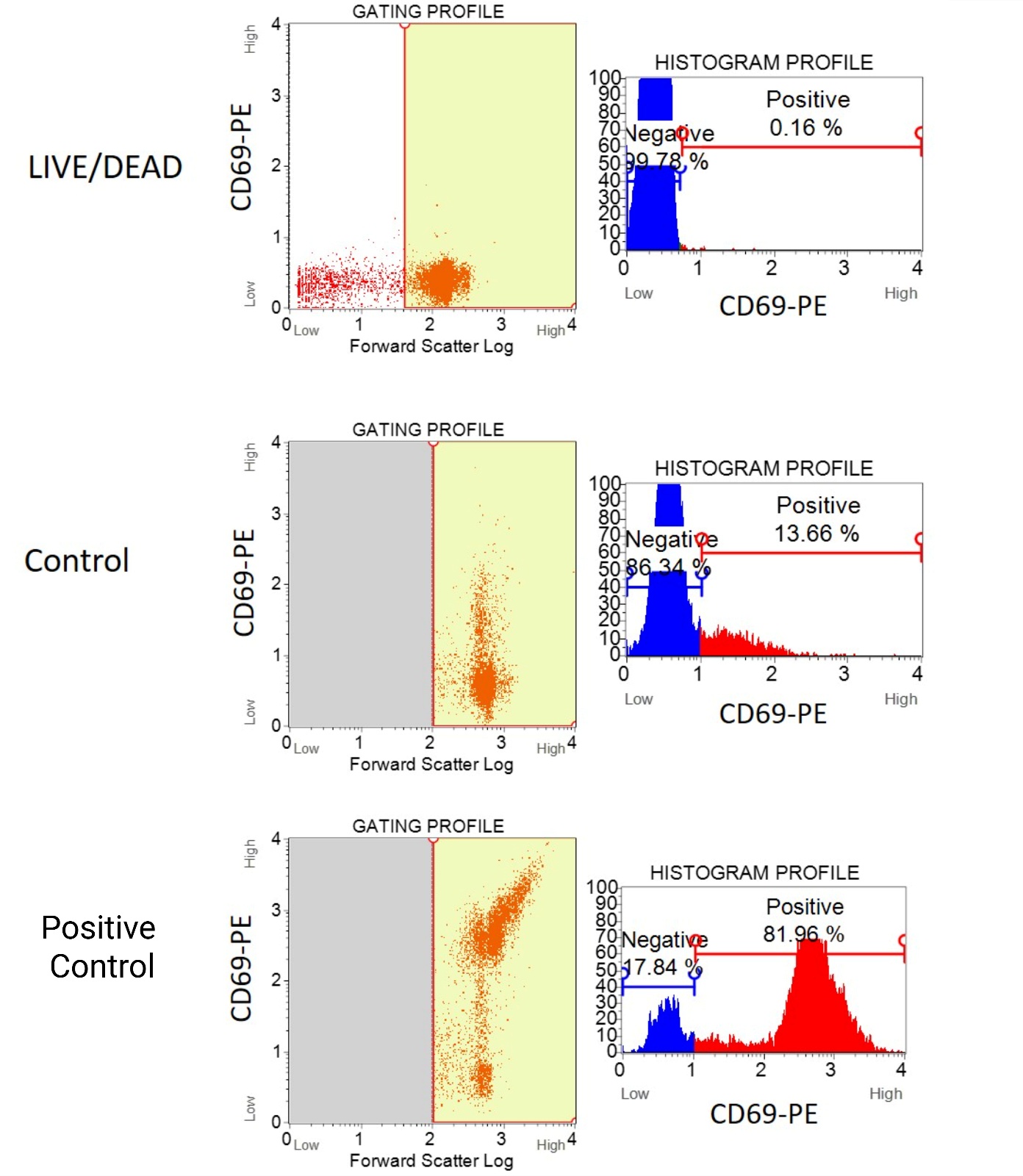

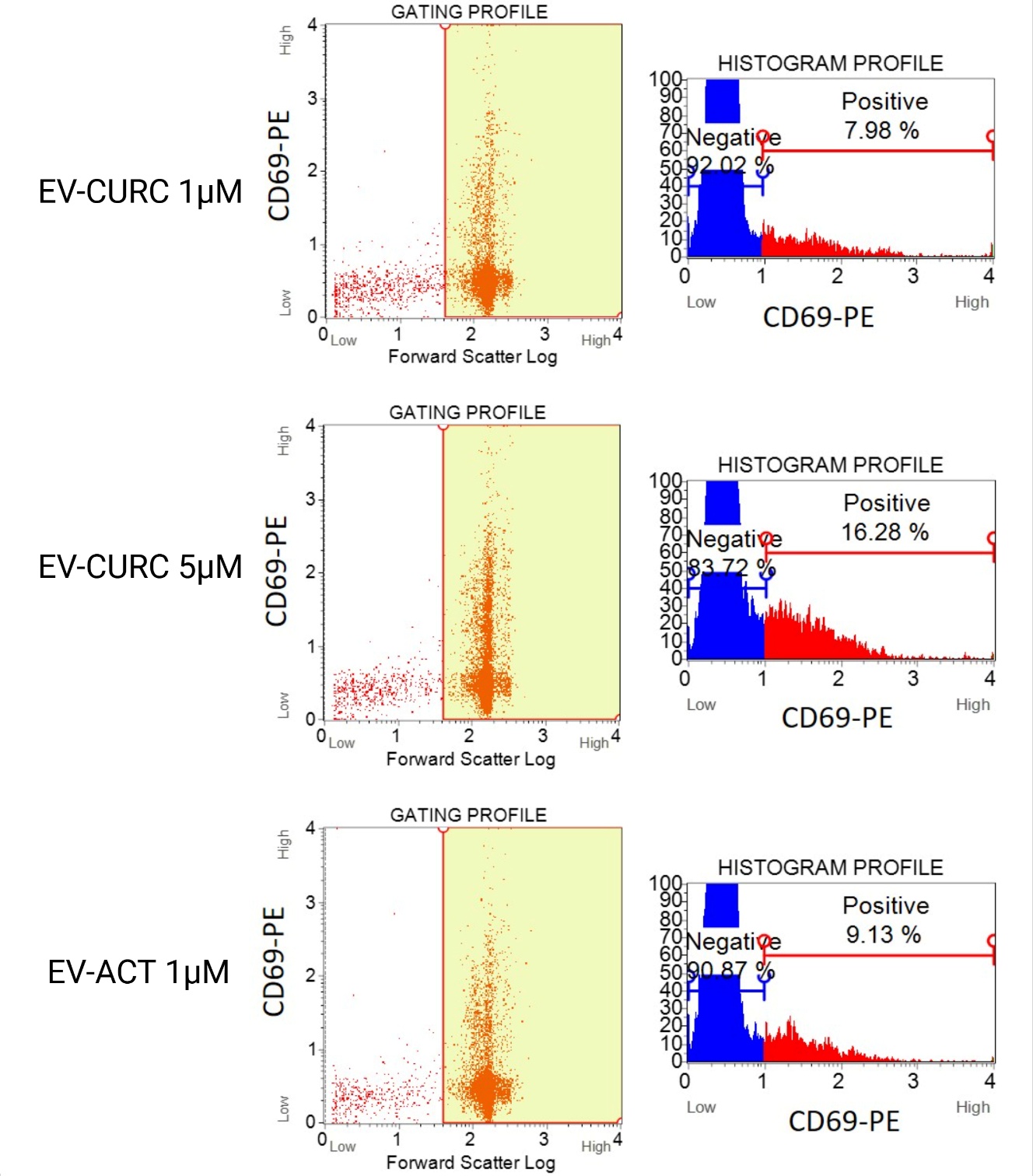

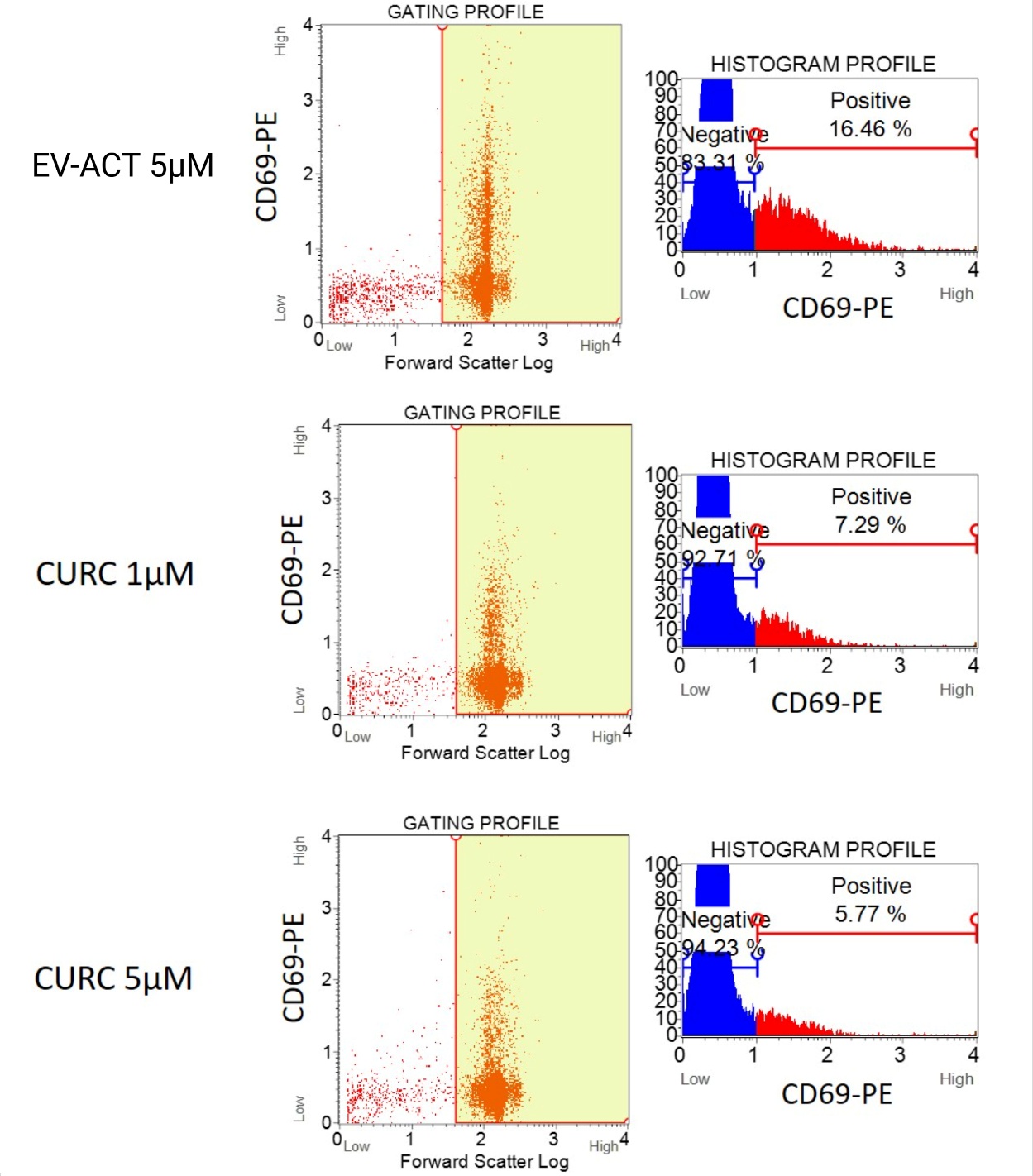

**Supplementary Figure S8**. Representative flow cytometry plots showing viability staining and CD69 expression in CD8^+^ T cells after 24h treatment with EV-CURC. The plots show gating strategy after staining with anti-CD69-PE (Left) and the corresponding histograms (Right).

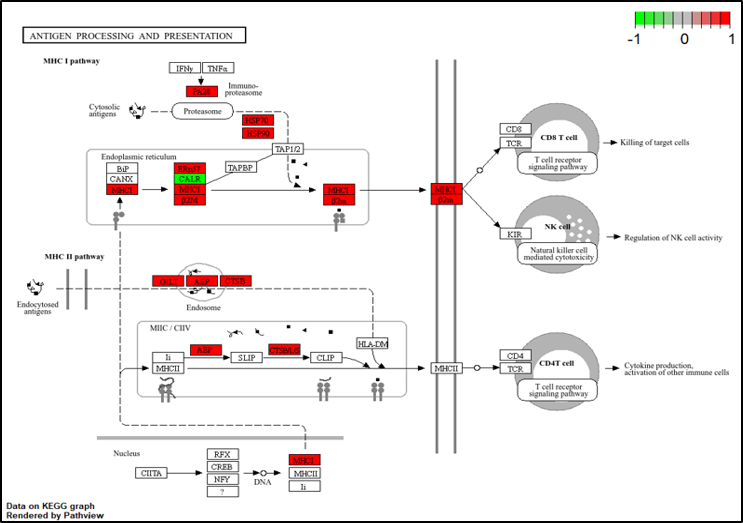

**Supplementary Figure S9.** Differentially abundant proteins mapped onto the Antigen processing and presentation KEGG pathway (mmu04612). Protein abundances were quantified by LC-MS/MS and were filtered (≥2 unique peptides). Colors indicate significant log2 fold change differentially abundant proteins involved in this pathway that are enriched in EV-ACT compared to EV-NA (red(0-1)), involved in antigen processing and presentation to CD8^+^ T cells, including proteins such as HSP70, HSP90, MHC-I/β2M complex. Red = dataset-enriched proteins; green = dataset decreased proteins; white = proteins that are not highlighted in our dataset (not-significant); grey = structural/contextual elements of the pathway that are not protein boxes.

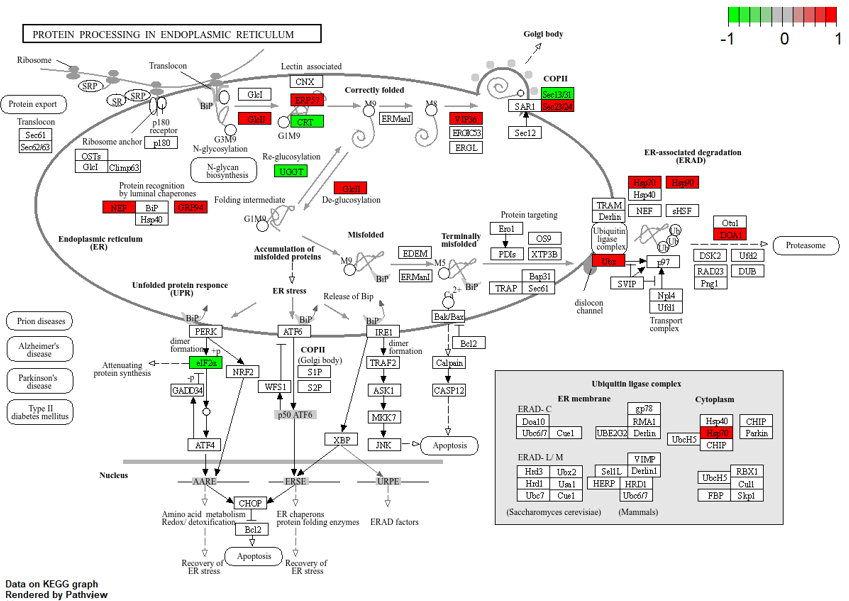

**Supplementary Figure S10.** Differentially abundant proteins mapped onto the Protein processing in endoplasmic reticulum KEGG pathway (map04141). Protein abundances were quantified by LC-MS/MS and were filtered (≥2 unique peptides). Colors indicate significant log2 fold change differentially abundant proteins involved in this pathway that are enriched in EV-ACT compared to EV-NA (red(0-1)), to highlight proteomic changes induced by CpG activation and TRP-2 peptide-pulsing in DC2.4 dendritic cells and to identify differences reflected in the EVs produced by these cells. Red = dataset-enriched proteins; green = dataset decreased proteins; white = proteins that are not highlighted in our dataset (not-significant); grey = structural/contextual elements of the pathway that are not protein boxes.

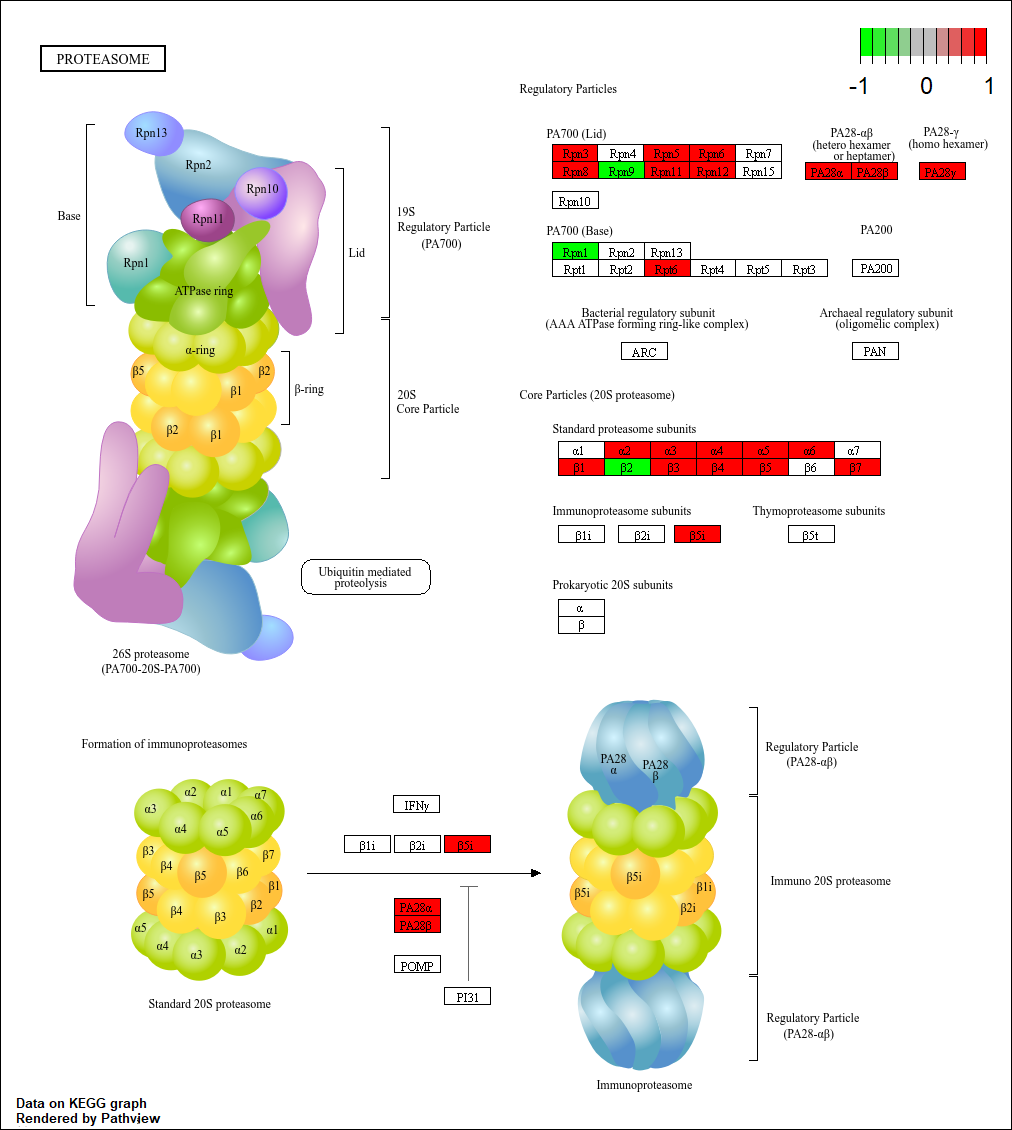

**Supplementary Figure S11.** Differentially abundant proteins mapped onto the Proteasome KEGG pathway (map03050). Protein abundances were quantified by LC-MS/MS and were filtered (≥2 unique peptides). Colors indicate significant log_2_ fold change differentially abundant proteins involved in this pathway that are enriched in the EV-ACT proteome compared to EV-NA (red(0-1)), with most of the proteins representing standard proteasome subunits, thereby indicating a cellular shift towards immune presentation. Red = dataset-enriched proteins; green = dataset decreased proteins; white = proteins that are not highlighted in our dataset (not-significant).

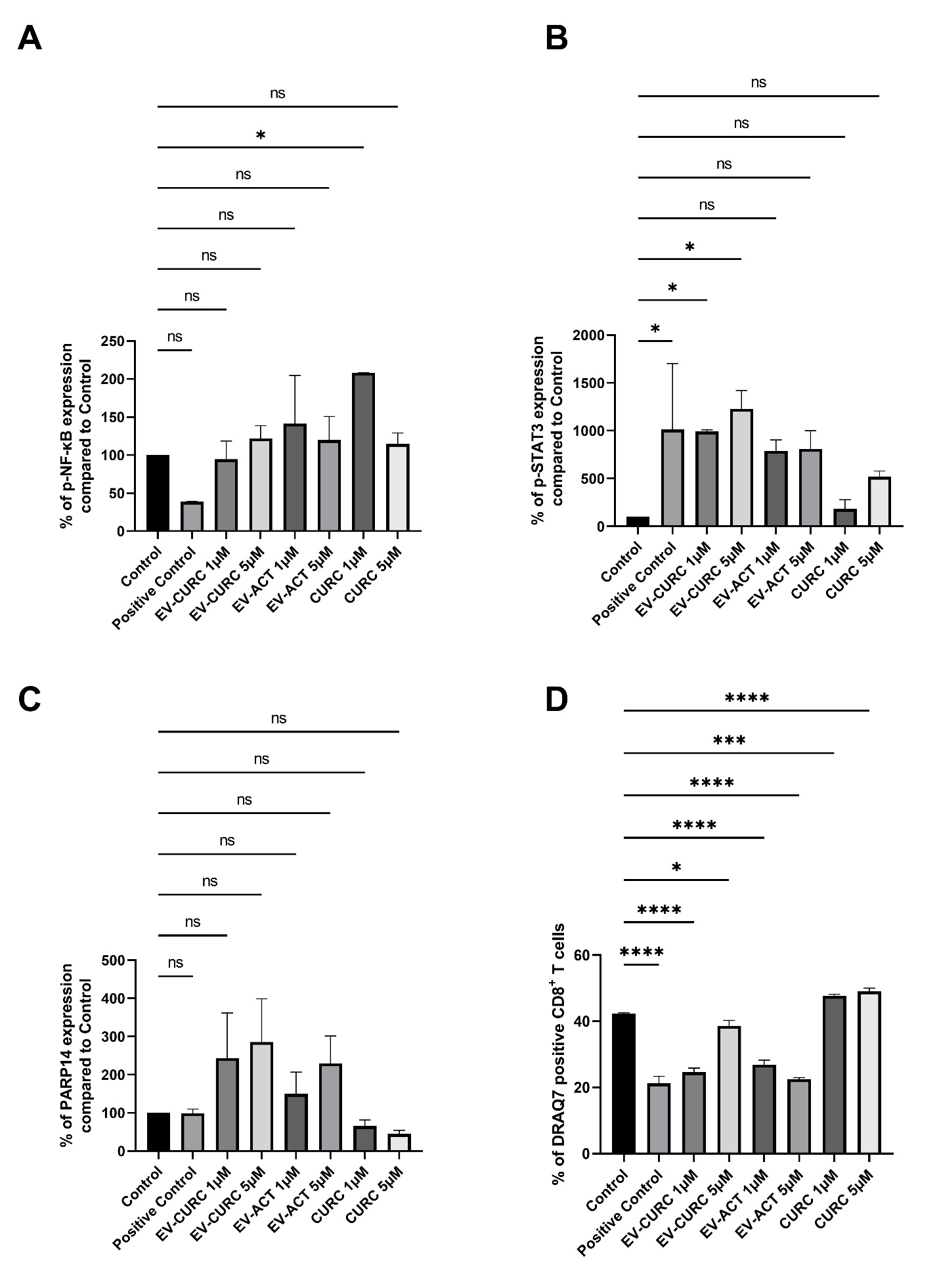
**Supplementary Figure S12.** Quantification of relative protein expression of p-NF-κB (**A**), p-STAT3 (**B**), PARP14 (**C**) and cell viability (**D**) in CD8^+^ T cells at 24 h. (n = 2 technical replicates). Statistical significance was analyzed by One Way ANOVA with Dunnett’s multiple comparisons test, data is expressed mean ± S.D.; ns – not significant; P > .05; ^*^, P < .05; ^**^, P< .01; ^***^, P < .001; ^****^, P <.0001.
